# Atomic structure of Hsp90:Cdc37:Cdk4 reveals Hsp90 regulates kinase via dramatic unfolding

**DOI:** 10.1101/040907

**Authors:** Kliment A. Verba, Ray Yu-Ruei Wang, Akihiko Arakawa, Yanxin Liu, Mikako Shirouzu, Shigeyuki Yokoyama, David A. Agard

## Abstract

The Hsp90 molecular chaperone and its Cdc37 co-chaperone help stabilize and activate over half of the human kinome. However, neither the mechanism by which these chaperones assist their client kinases nor why some kinases are addicted to Hsp90 while closely related family members are independent is known. Missing has been any structural understanding of these interactions, with no full-length structures of human Hsp90, Cdc37 or either of these proteins with a kinase. Here we report a 3.9Å cryoEM structure of the Hsp90:Cdc37:Cdk4 kinase complex. Cdk4 is in a novel conformation, with its two lobes completely separated. Cdc37 mimics part of the kinase N-lobe, stabilizing an open kinase conformation by wedging itself between the two lobes. Finally, Hsp90 clamps around the unfolded kinase β5 strand and interacts with exposed N-and C-lobe interfaces, safely trapping the kinase in an unfolded state. Based on this novel structure and extensive previous data, we propose unifying conceptual and mechanistic models of chaperone-kinase interactions.

**One Sentence Summary:** The first structure of a chaperone:kinase complex reveals that the Hsp90 system modulates and stabilizes kinases via a functionally relevant, unfolded open state.

## Main Text

The human kinome is responsible for regulating about a third of all proteins via phosphorylation(*1*). Proper regulation of this process is of paramount importance, as misregulated kinase activity can lead to cell death and disease(*2*). In keeping with their importance, kinase activity can be sensitively modulated by multiple allosteric inputs. Thus, kinase domains are organized such that dispersed small structural changes due to binding of regulatory domains/proteins or phosphorylation, can significantly alter kinase activity. Examples of such regulator interactions abound, via SH2/SH3 domains for Src family kinases, dimerization for EGFR or Raf family kinases, and cyclin regulation for Cdks being well characterized examples(*3*).

Beyond these specific regulators, the Hsp90 molecular chaperone, a member of the general cellular protein folding machinery, also plays a fundamental role in the regulation of many kinases(*4*). While usually chaperones facilitate the early steps of protein folding, Hsp90 uniquely functions late in the folding process to help both fold and activate a privileged set of protein “clients” (~10% of the proteome)(*5*). Notably ~60% of the human kinome interacts with Hsp90 with the assistance of its kinase specific cochaperone Cdc37(*6*). Pharmocologic inhibition of Hsp90 leads to rapid ubiquitinylation and degradation of client kinases. As many Hsp90/Cdc37-dependent kinases are key oncoproteins, (vSrc, bRafV600E, Her2, etc.) several Hsp90 inhibitors are undergoing clinical trials as cancer therapeutics(*7*).

Hsp90 is a well conserved, but highly dynamic molecular machine. Each monomer within the Hsp90 dimer has three structural domains: a C-terminal domain (CTD) responsible for dimerization; a middle domain (MD) implicated in client binding; and the N-terminal domain (NTD) binds ATP. Without nucleotide Hsp90 mostly populates a variety of “open” states, whereas nucleotide binding promotes formation of a closed state in which the NTDs also dimerize, followed by hydrolysis(*8, 9*). The rates of closure and hydrolysis are homologue specific, with human cytosolic Hsp90s almost always open, while yeast Hsp90 adopts a fully closed state, suggesting a species specific optimization(*10*). Towards the end of the NTD is a highly charged region (“charged linker”) that varies wildly in length and composition between species. The function and the structure of the charged linker are unclear, but deletion can impact function(*11*).

By contrast, Cdc37 is considerably less well studied. The monomeric protein can also be divided into three domains: an N-terminal domain of unknown structure that interacts with kinases, a globular middle domain which interacts with Hsp90 and an extended C terminal domain, of unknown function(*12*). Although there is a cocrystal structure of the Cdc37 middle/C domains (Cdc37 M/C) bound to the Hsp90-NTD(*13*), there is evidence that Cdc37 may also interact with the MD of Hsp90(*14*). Phosphorylation of Cdc37 serine 13 plays an important role, providing stabilizing interactions *in vitro*(*15*) and being functionally necessary *in vivo*(*16*).

Although there is a wealth of *in vivo* data, a physical understanding of how Hsp90 and Cdc37 facilitate kinase function is lacking. Equally unclear is why some kinases are strongly Hsp90-dependent whereas other very closely related kinases are Hsp90-independent. Despite numerous attempts to identify a consistent motif responsible for Hsp90 interaction, the only general trend that has emerged is that client kinases appear to be less thermally stable than non-clients(*6*). In support of this, binding of kinase inhibitors or allosteric regulators reduce Hsp90 interactions(*17, 18*). While reasonable that less stable kinases might depend on Hsp90, why this happens and what Hsp90 is recognizing has yet to be established. Despite its obvious value, obtaining a crystal structure of an Hsp90:Cdc37:kinase complex has been unsuccessful due to the dynamic nature of Hsp90-client interactions, and challenges in reconstituting the complex. Encouraged by previous negative stain EM studies(*19*) and the stunning recent advances in cryoEM detectors and processing methodologies that together make analysis of such a small, asymmetric complex feasible(*20*), we undertook cryoEM studies of the human 240KDa Hsp90:Cdc37:Cdk4 kinase complex.

### Complex formation and cryoEM structural analysis

Human homologues of all three proteins were co-expressed in Sf9 insect cells. When trapped with molybdate, a stable ternary complex formed capable of surviving a rigorous dual-tag purification (Fig S1). Although the mode of molybdate action is unknown, it affects Hsp90’s hydrolysis rate and helps stabilize Hsp90-client interactions(*21*). Simply mixing components individually purified from insect cells failed to form any detectable complex; therefore either post translational modifications specific to the complex or other components (like Hsp40/Hsp70) are required. Despite significant effort to optimize conditions and image processing, preferential particle orientation and conformational heterogeneity limited initial reconstructions from data collected in house to about 6-8Å resolution. To move forward a much larger data set was collected at NRAMM (See methods). Data quality was verified as 2D classification of ~800,000 initially picked particles yielded classes with visible secondary structure (Fig S2). 3D classification and refinement resulted in a 4Å map (Fig S3 and Table S1), with most of Hsp90 being better (3.5Å) while the lowest resolution regions were around 6Å (Fig 1A).

**Fig. 1.**
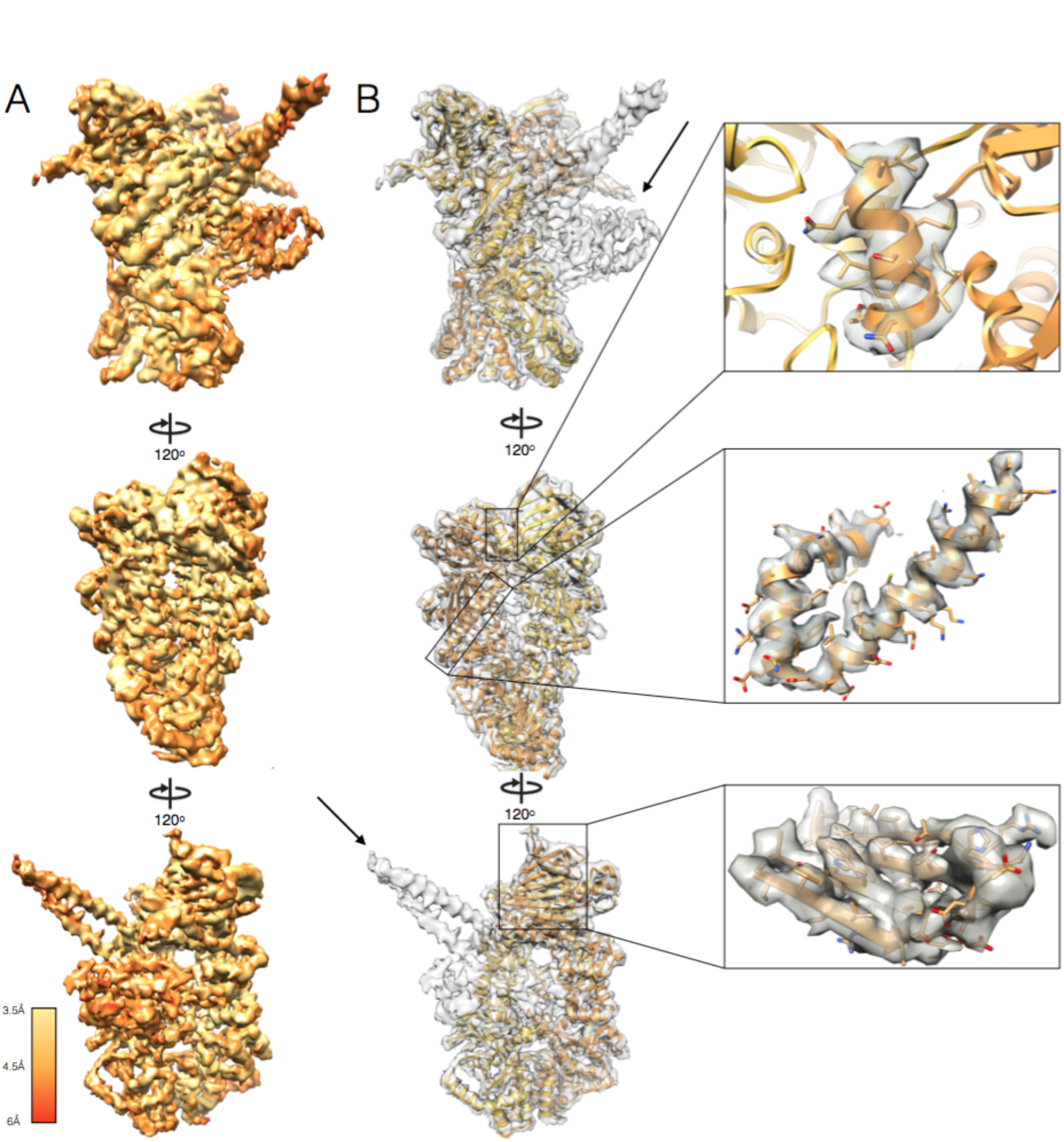
The 4Å map of Hsp90/Cdc37/Cdk4. (A) Density map colored by resolution. (B) hHsp90β model built into the density map, with different monomers colored shades of orange. Inserts show high resolution features. Arrows show density un-accounted for by Hsp90.

### Hsp90 is in a closed conformation in the complex

Refining the 4Å map with a tighter mask around Hsp90 generated a higher resolution, 3.9Å density for Hsp90 and neighboring regions (Fig. 1B inserts)(See methods). This map was of sufficient quality to accurately build/refine an atomic model of human Hsp90b based on a homology model derived from yeast Hsp90 (Fig 1B and S4). Surprisingly, reconstructions with an Hsp90 charged linker deletion never yielded good results, likely due to increased heterogeneity. Hsp90 adopts a symmetrical (RMSD between monomers=0.82Å), closed conformation, closely resembling the yHsp90 closed state(*22*). This was unexpected, as without crosslinker, closed hHsp90 had never been observed. While the molybdate added during purification may contribute, it is likely a consequence of the ternary complex. In our map, which was refined without symmetry, at very high thresholds there is a strong symmetric density at the γ-phosphate location for nucleotide binding sites at both NTDs of Hsp90 (Fig S5). This suggests that either the complex traps Hsp90 in an ATP state, or, more likely, ADP-molybdate acts as a post hydrolysis transition state inhibitor, thereby helping stabilize the closed conformation. The hHsp90 model determined here is similar to the yeast Hsp90 (RMSD=1.59Å), with a small rotation in the CTD and correlated movements throughout (Movie S1). Additionally, a number of loops disordered in the crystal structure were ordered in our model, as they were interacting with the kinase. (Fig 2B) Also, this is the first structure with an intact charged linker. While the density in the reconstruction and 2D classes extends beyond the truncated regions seen in the crystal structure, the 3D density is weak suggesting a highly flexible structure (Fig S2).

**Fig. 2.**
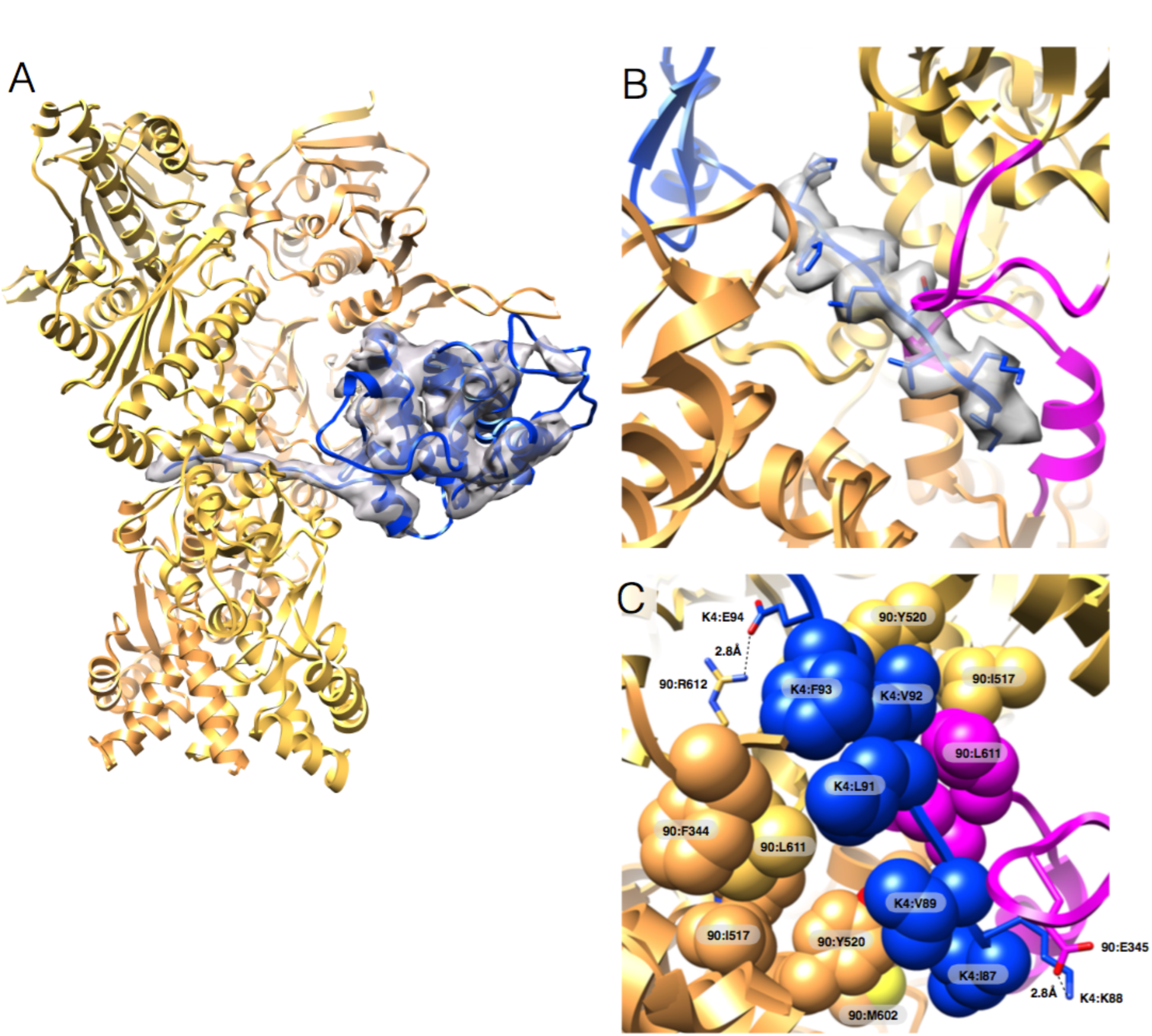
Cdk4 is unfolded when complexed with Hsp90 and Cdc37. (A) Cdk4 (K4) C-lobe (blue) fit into the map (B) The tubular density from high-resolution map through the lumen of Hsp90 is perfectly fit by an unfolded β5 sheet (in sticks) of Cdk4. In magenta are previously disordered client interacting loops on Hsp90. (C) Cdk4/Hsp90 interface with hydrophobic residues in spheres, salt bridges in sticks.

### The N-lobe of Cdk4 is significantly unfolded and threads through Hsp90

Two regions of the 4Å map cannot be accounted for by Hsp90: a long coiled-coil like protrusion and a globular density on one side of Hsp90 (arrows, Fig 1B). While we initially suspected that these corresponded to the Cdc37M/C (globular domain with long helix), as our map improved the globular domain fit became worse. To obtain an unbiased fit, an exhaustive search was performed against all unique protein folds (See methods). Disregarding algorithmic errors, the top scoring hits were all variations on the kinase C-lobe, with Cdc37M/C scoring considerably worse (Fig S6). Based on this, the Cdk4 C-lobe was placed into the density(*23*), resulting in a high quality fit (Fig 2A). By contrast, no suitable density exists for the kinase N-lobe. Moreover, a folded N-lobe would sterically clash with Hsp90. Remarkably, tracing the kinase density from the C-lobe towards the N-terminus, there was a clear tubular region going through the lumen of Hsp90 (Fig 2A, 2B). Thus, a drastically altered conformation of the kinase N-lobe is being stabilized by Hsp90. (Movie S2) Threading the Cdk4 sequence into this density, reveals that β4-β5 strands have been ripped apart and instead b5 interacts with a previously mapped general Hsp90 client binding site via extensive hydrophobic interactions(*24*) and two salt bridges (Fig 2C). Potentially due to these interactions, there is a rotation of Hsp90 CTD towards the kinase (as compared to yHsp90 structure) at MD:CTD interface, which was identified as asymmetric in previously published work(*25*) (Movie S1). Altogether this suggests that MD:CTD interface may be used to communicate between client and Hsp90’s hydrolysis state. Unfortunately, in the highest resolution map no density was visible for kinase residues N-terminal of the β4 strand.

### Cdc37 is split into two domains and wraps around Hsp90

While there is no available 3D structural information for the Cdc37 N-terminal domain, sequence analysis (MARCOIL) predicts significant coiled coil structure. Supported by our observations of high helical content by CD and NMR (Fig S7), this provides a good candidate for the non-globular map density (Fig 1B). That density, transitions from helical to strand like, wrapping around the Hsp90MD, adding an additional β-strand to the 1AC β-sheet (2CG9 nomenclature). Unfortunately, as with the kinase, this b-strand does not connect to any density on the other side, thwarting a complete fit of Cdc37. Showing the unique power of single particle cryoEM, local rounds of 3D classification yielded a 7Å map (Fig S8) showing clear globular density that connected through the β-strand to the coiled-coil region (Fig 3A). This new density was unambiguously fit by the crystallized Cdc37M/C fragment. Using a combination of the two maps, we were able to fit Cdc37M/C residues 148-260 and de-novo build residues 1-147 of human Cdc37 (Fig 3A) (See methods). Although further classification revealed some density beyond residue 260, it was too weak to model with confidence.

**Fig. 3.**
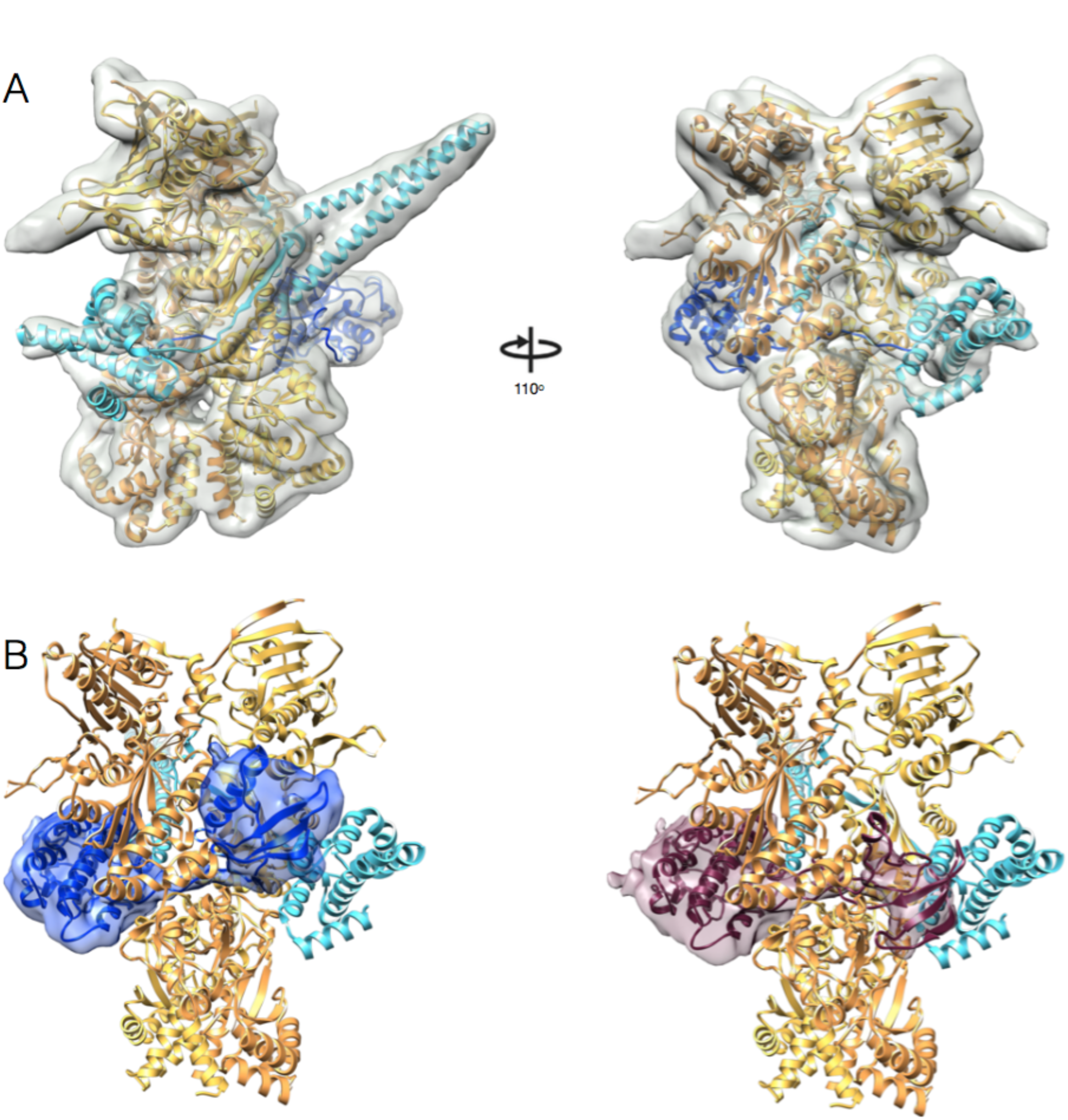
Rounds of focused 3D classification yield distinct densities for Cdc37 and the Cdk4 N-lobe. (A) One of the new classes has clear density for the Cdc37 (37) M/C fragment crystal structure. In teal is our complete Cdc37 model (residues 1-260). Note the b-strand wrapping around the outside of Hsp90 connecting the two major Cdc37 domains. (B) Two additional classes show new density for the missing Cdk4 N-lobe. The classes minus the Hsp90/Cdc37 density, in blue and maroon, highlight the two new Cdk4 N lobe conformations. Fitted kinase models are in ribbons.

### Locating the remainder of the kinase N-lobe and verification

The local 3D classification also revealed the remainder of the kinase N-lobe (Fig 3B and Movie S3). In two of the classes, distinct globular densities were visible that connected to the tube of density derived from the unfolded Cdk4 N-lobe. At the same time, the density for the middle domain of Cdc37 was either very weak or absent, suggesting an anti-correlation between Cdc37M/C:Hsp90 interactions and kinase N-lobe:Hsp90 interactions. We were able to roughly fit the rest of the Cdk4N lobe into these densities, allowing us to build a complete model of Hsp90:Cdc37:Cdk4 complex (Fig 4 and Movie S4). Given the unusual split domain conformation for both the kinase and cochaperone, it was important to verify our structural assignments. Towards that end, we covalently tagged Cdc37 at its N terminus and Cdk4 at its C terminus with T4 lysozyme which should be readily visible even in moderate resolution maps. The two different complexes were expressed and purified from yeast, followed by cryoEM reconstruction of each. These two structures clearly show lysozyme density consistent with our map interpretation (Fig S9).

**Fig. 4.**
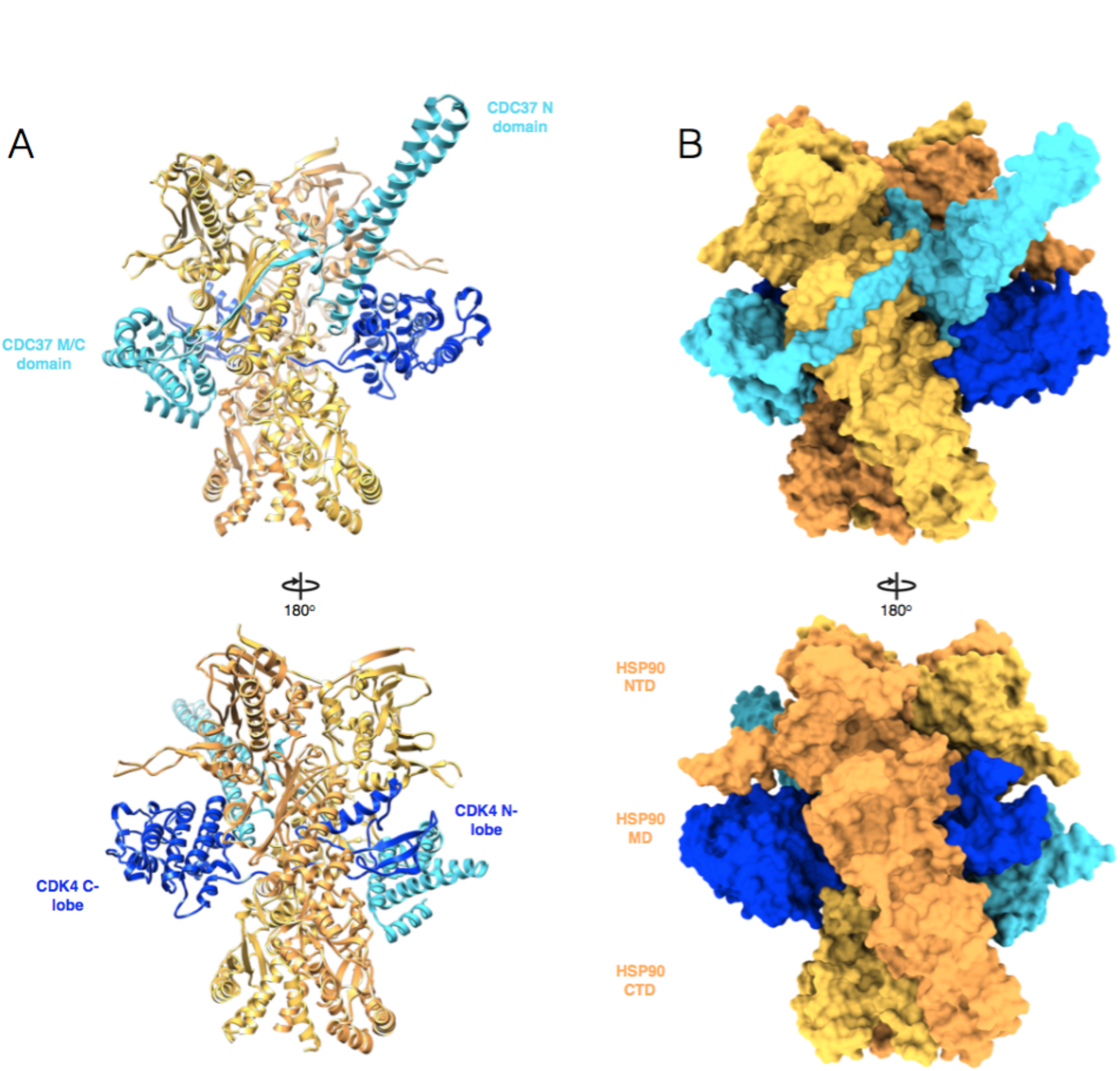
Hsp90, Cdc37, and Cdk4 are intricately interwoven in the complex. Two views of the complete model, showing ribbon in (A) and surface in (B)

### Cdc37 precisely mimics a conserved feature of kinase N-lobe-C-lobe interactions and makes unique interactions with Hsp90

There are a number of striking features from the resulting model (Fig 4) that explain a wealth of accumulated, seemingly contradictory observations. The Cdc37NTD forms a long coiled-coil with a leucine zipper like motif. The N-terminus of this coiled-coil interacts with Cdk4 through extensive hydrophobic and hydrogen bonding interactions, burying 725 Å^2^ (Table S2) in surface area, and mimicking interactions that the kinase αC-β4 loop normally makes with the C-lobe (Fig 5A). More strikingly, the following loop in Cdc37, overlays perfectly with the αC-β4 loop of multiple kinases, packing tightly against the kinase C lobe (Fig 5A, bottom insert), explaining why this Cdc37 region is so conserved (Fig. S10). Thus Cdc37 binds to the kinase C-lobe by mimicking interactions the N-lobe would normally have made, stabilizing separation of the two kinase lobes.

**Fig. 5.**
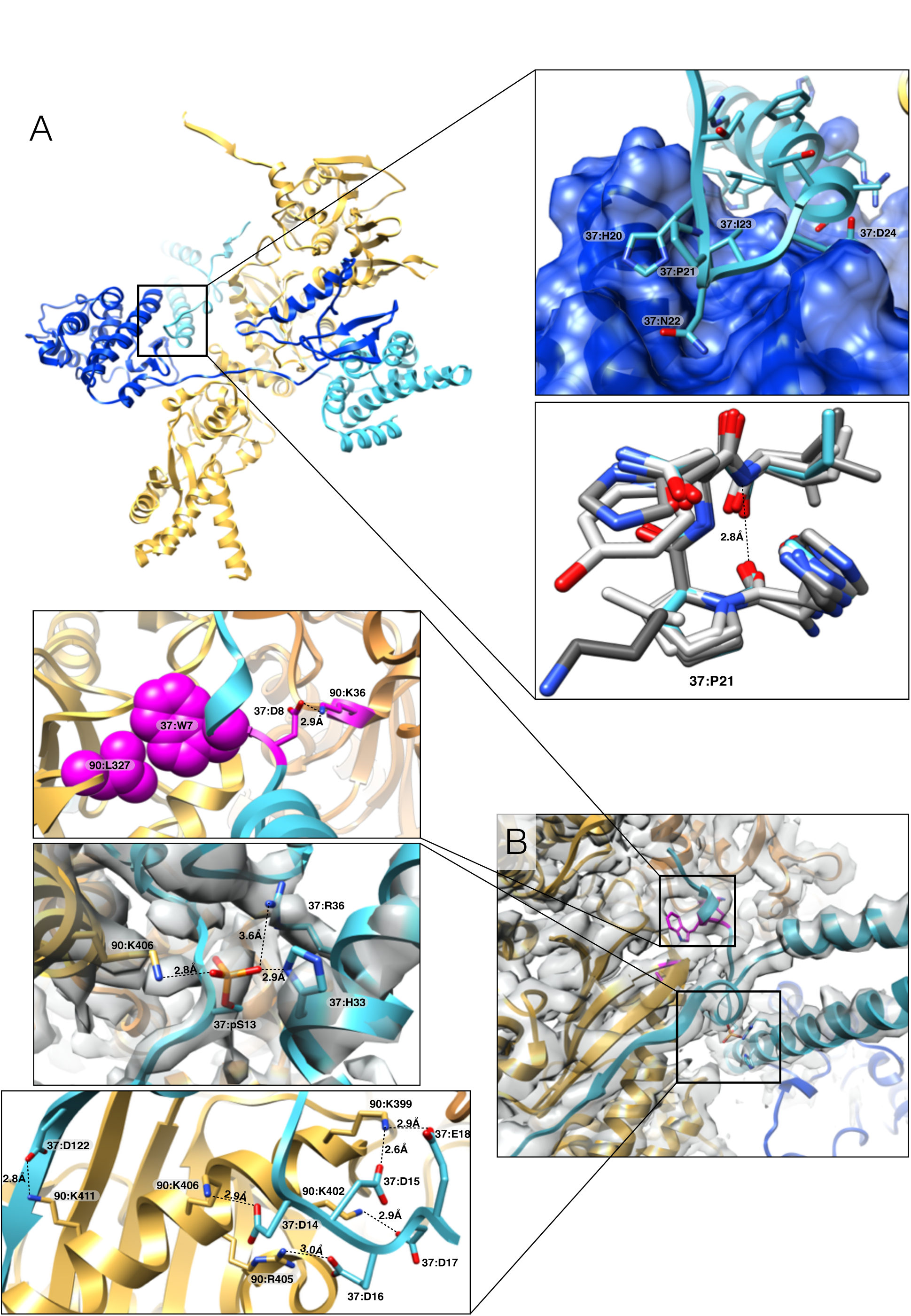
High-resolution details of Cdc37 interactions with Hsp90 and Cdk4. (A) Overall arrangement of Hsp90/Cdc37/Cdk4 (one Hsp90 monomer removed for clarity). The insets highlight Cdc37/Cdk4 interaction features: top - Cdc37/Cdk4 interact via hydrophobic interactions and backbone hydrogen bonds, with perfect shape complementarity, bottom-overlay of Cdc37’s conserved HPN motif (teal) perfectly mimicking type I β-turn of αC-β4 loop of 6 different kinases (shades of gray). (PDB codes: 3g33, 2itp, 1qmz, 3pp0, 1jnk, 4fk3) (B) Zooming in on Hsp90/Cdc37 interactions. Top insert: Cdc37/Hsp90 interactions mimic p23/Hsp90 interactions identified previously (magenta). Middle insert: phospho-Ser13 stabilizes local Cdc37 structure through interactions with conserved R36 and H33 and also interacts with Hsp90 at K406. Bottom insert: six salt bridges stabilizing Hsp90/Cdc37 interactions.

The interactions between Cdc37 and Hsp90 are significantly different from those seen in the crystal structure of fragments of each protein. Instead of binding to an Hsp90NTD surface that would only be accessible in the open state, in the Hsp90 closed state the Cdc37M/C interacts with Hsp90MD, although these interactions are fairly limited (only about 340Å^2^ out of 2650Å^2^ total buried surface area) (Fig 4). More extensive are interactions from Cdc37 residues 120-129, which bind to the Hsp90 MD and contribute an additional β-strand (Fig 5B). Additionally part of the Cdc37NTD binds to a new site on the closed Hsp90NTD, somewhat mimicking interactions seen between p23 and Hsp90 (Fig 5B, top insert). There is also a network of ionic interactions between Cdc37NTD and Hsp90MD, which explains previously identified salt sensitivity of Cdc37/Hsp90 binding (Fig 5B, bottom insert). Our structure thus helps explain recent data that *C. elegans* Cdc37 interacts with the middle domain of Hsp90(*26*), rather than the NTD as observed in previous human domain binding studies. Instead of assuming the *C. elegans* is somehow unique or that the interactions observed with isolated domains were incorrect, we propose that Cdc37/kinase first binds Hsp90 in its open conformation much as in the crystal structure, and then rearranges to the site on the middle domain upon Hsp90 closure, as seen in our structure (see model, below).

### Conservation of Cdc37 interactions and phosphorylation

The residues comprising Cdc37/Hsp90NTD and Cdc37/kinase interaction surfaces are extremely well conserved (Fig S10), further validating the significance of the interactions observed here. Fungal Cdc37s have a long insertion, that by being located in the loop between the two long coiled-coil α-helices, would not disrupt any interactions seen in the human complex. This suggests that this prominent structure may serve some additional function or regulatory role, optimized by extension in fungi.

Phosphorylation at the perfectly conserved Cdc37:S13 is important for kinases to function, and was thought to be directly involved in kinase binding (*15,16*). In our structure there is density for the phosphorylated-serine13, which forms a salt bridge with Cdc37:R36 and Cdc37:H33, stabilizing the very N-terminus of the coiled coil (Fig 5B middle insert). This provides a molecular rationale for previous observations of a Cdc37 conformational change upon phosphorylation(*27*). Of note, this phosphate also contributes to the overall electrostatic nature of Hsp90-Cdc37 interactions, forming a salt bridge with Hsp90:K406.

### Hsp90/Cdc37 capture and stabilize structural transitions within the kinase

In our structure, Cdk4 assumes a conformation drastically different than previously seen for any kinase structure. The kinase hinge region and β4-β5 sheets are completely unfolded, with the N lobe and C lobe being pried apart and stabilized by new interactions with Hsp90 and Cdc37. Based on this, we propose a model where it is not a specific binding sequence, but the propensity for the kinase to access an N-lobe/C-lobe separated open state (unfold), that determines whether a particular kinase will be a client. In agreement with this, the β3-αC loop of Cdk4 (strong client) has seven glycines versus one/two glycines in the same loop of Cdk2 and Cdk6 (non client and weak client respectively). Mutating this loop in Cdk4 to the Cdk6 sequence stabilizes the protein(*28*). In a similar manner, a single point mutation derived from EGFR (non-client) in the αC-β4 loop was able to abrogate the interactions of HER2 (strong client) with, and dependence on, Hsp90/Cdc37(*29*). This point mutation would potentially introduce a salt bridge stabilizing the kinase loop (Fig S11), and hence the association between the kinase lobes.

Our model also explains why sequence mutations that alter Hsp90 interactions have often been kinase idiosyncratic, providing no general understanding. Indeed, the remarkable allostery so important for kinase regulation allows distant mutations to destabilize N-lobe/C-lobe interactions. It is also consistent with observations that some kinases only depend on Hsp90/Cdc37 during their initial folding whereas closely related family members remain addicted to Hsp90/Cdc37 throughout their functional lifetimes. Furthermore, this argues that the kinase seen in our structure may be a trapped form of an on pathway folding intermediate, only visualizable due to its stabilization by Hsp90/Cdc37 (Fig 6). Most if not all kinases would be expected to go through such a state during their initial folding, but most would then be strongly stabilized in the folded state via either intra or intermolecular interactions (cyclin, SH2/SH3, dimerization, etc.). By contrast, client kinases are not so stabilized and would more readily sample this state.

**Fig. 6.**
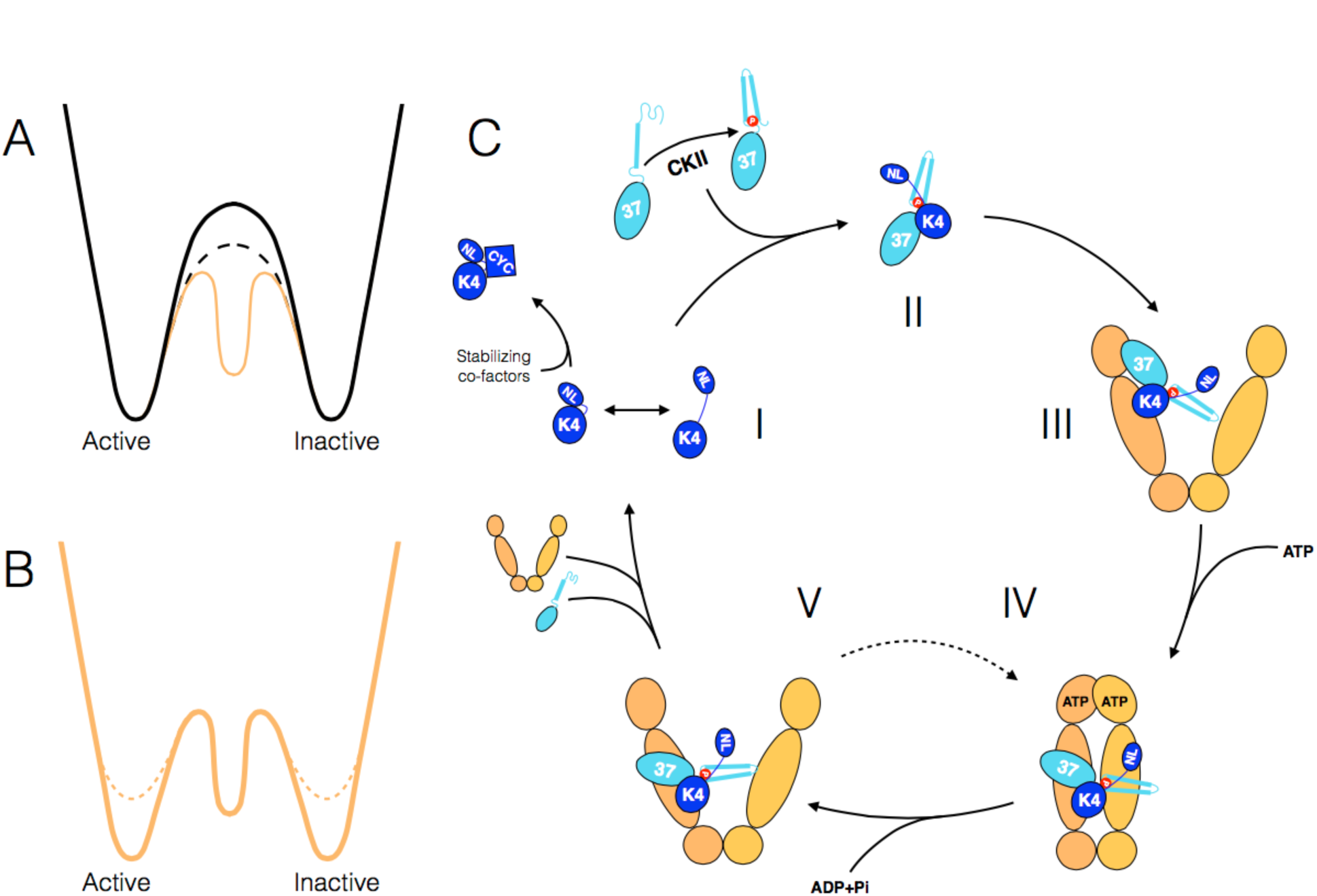
Conceptual model for linkage between kinase folding and activation and proposed model for the Hsp90:Cdc37:kinase cycle. (A) Transitioning between states through an unfolded intermediate (dashed line) has a lower energy barrier than through rigid body motion (solid line). The Hsp90:Cdc37 stabilizes such an unfolded intermediate (orange solid line) (B). By comparison with non-clients (solid line), the active and inactive states of client kinases (dashed line) are de-stabilized. (C) Speculative model for an Hsp90:Cdc37:Kinase cycle. (I) The kinase domain transiently samples an open state. Interactions with co-factors (like cyclins, SH2/SH3 domains, etc.) stabilizes the kinase native state, disfavoring the open state. (NL-N-lobe, CYC-Cyclin) CKII phosphorylated Cdc37 captures the open state by binding the kinase C-lobe (II). Cdc37/kinase then binds to open Hsp90 (III). Hsp90 binds to ATP and closes upon the unfolded part of the kinase. Cdc37 migrates down, resulting in the structure described here (IV). Upon hydrolysis of ATP, Hsp90 opens with Cdc37/Cdk4 still bound, giving a chance for the kinase to fold (V). If it folds, it displaces Cdc37 and leaves the complex. If however it fails to displace Cdc37, then Hsp90 is able to re-bind ATP and go back to state IV, repeating the process. At some point during this cycle, PP5 phosphatase is recruited to the complex to de-phosphorylate Cdc37.

### Linking kinase unfolding to assembly and activity

Given such a model, why might it be beneficial for a large percentage of human kinases to significantly populate a folding intermediate even when mature? One explanation could be that given the importance of kinases for so many aspects of cellular and organismal regulation, diversifying functional properties likely arose via destabilizing mutations, which were then “buffered” by Hsp90/Cdc37. Since there was little cost, these mutations could then become fixed in the population. Much data already supports the ability of Hsp90 to buffer phenotypic variation (*30*). Similarly, many cancers become addicted to high levels Hsp90/Cdc37 as their driving oncogenic kinases need chaperone stabilization(*31*). However, if Hsp90s were only important for their initial evolution, it might be expected that client kinases would eventually be stabilized by secondary mutations. An alternative model, which we favor, is that being able to safely populate such an open folding intermediate itself has direct functional/regulatory benefit.

Interestingly, computational efforts have suggested a connection between folding and kinase activity(*32*, *33*). The concept is that the most favorable way to transition between the inactive and active states is to transition through a more open, unfolded state rather than through a more classical rigid-body transition. Although the opening transition seen in simulations is far more subtle than that observed here, we suggest that the concept still applies (Fig 6A). The generally lower stability of client kinases would lead to enhanced sampling of the open state, thereby encouraging chaperone binding (Fig 6B). Chaperone stabilization of a kinase open state could increase the overall rates of interconversion and/or protect a potentially vulnerable state from either aggregation or recognition by the ubiquitinylation machinery. Moreover, the open state could be the preferred substrate for adding/removing post-translational modifications as well as for facilitating a dynamic equilibrium with critical stabilizing interactions.

### Life cycle of kinase:Hsp90:Cdc37 interactions

Our structure also provides a basis for speculation on how the observed state might arise and mature (Fig 6C). Entry into the cycle would be dependent on the ability of Ser13 phosphorylated Cdc37 to outcompete the native kinase αC-β4 loop for interaction with the C lobe. Once bound, Cdc37 could deliver the kinase to an open Hsp90, perhaps via previously published crystal structure contacts. Whether this step also requires assistance from Hsp70/Hsp40 as with the glucocorticoid receptor is as yet unclear(*34*), but the inability to directly form the complex from components and Chk1 reconstitution experiments(*35*) are suggestive. Concomitant with ATP-dependent Hsp90 closure, Cdc37 would migrate to the middle domain using an alternative surface, forming the structure seen here (Fig 4). By opening up, the long coiled coil domain could potentially allow Cdc37/kinase to stay connected with Hsp90 during multiple ATPase open/closure cycles, with the Hsp90 ATPase playing a timing role. Upon Hsp90 opening, the kinase N-lobe would have a finite time to fold and outcompete Cdc37’s N-terminal domain for the C-lobe, before Hsp90 would bind another ATP and close again. In a non-client kinase (ie, during initial folding), the N-lobe would likely efficiently outcompete Cdc37, promoting dissociation from Hsp90/Cdc37 and subsequent Hsp90 independence. In such a model, Cdc37 provides a quality control checkpoint, where only upon kinase folding will it be outcompeted. While this captures the essence of the available data, other models are possible.

A client kinase such as Cdk4 may spend a substantial amount of time in the protected yet inactive Hsp90/Cdc37 bound state, waiting for cyclin D. Once available, a high cyclin D concentration would provide sufficient stabilization energy for folded Cdk4 to outcompete Cdc37, freeing the kinase domain. Alternatively, if the Hsp90/Cdc37/kinase persists too long in the Hsp90 open state (very unstable kinase or Hsp90 inhibitor added), then Hsp70 binding might be preferred as with GR, facilitating recruitment of the degradation machinery to the kinase.

Beyond revealing the kinase open state, our cryo EM reconstruction allowed us to build first atomic models for the human cytosolic Hsp90 and the kinase interacting N terminus of Cdc37. This is the first high-resolution structure of Hsp90 interacting with a client of any kind, and of full length Hsp90 interacting with Cdc37. Moreover, the ability to collect a large number of particles coupled with the capabilities of single electron counting detectors and 3D classification software, allowed us to visualize multiple conformations, providing a qualitative assessment for the dynamic nature of the complex. Overall, our structure has explained a number of often contradictory, biochemical observations and provided both mechanistic and conceptual models of Hsp90:kinase interactions, that can be tested in future experiments. It also indicates the potential of single particle cryoEM for exploring other challenging dynamic, asymmetric complexes at near atomic resolution.

## Materials and Methods

### Expression and purification of Hsp90/Cdc37/Cdk4 complex

#### Expression of the complex in Sf9 cells

Such prepared sample was used for all the reconstructions except T4 Lysozyme labeled complexes. The plasmids encoding the full-length human HSP90b and Cdc37 were kindly provided by Dr. Neil F Rebbe (The University of North Carolina at Chapel Hill) and Dr. Ernest Laue (The University of Cambridge), respectively. The cDNA clone for Cdk4 was purchased from Origine.

The DNA fragment of the full-length Hsp90b (residues 1-724) or the linker-deleted Hsp90b (residues 1-220-GGGG-274-724), in which the linker region (residues 221-273) were replaced by GGGG, was independently amplified by PCR and subcloned into the baculovirus transfer vector pFastBacHT (Thermo Fisher Scientific, USA) as a fusion with an N-terminal Flag tag and a TEV cleavage site. The DNA fragment of the Cdk4 (residues 1-303) for a fusion with an N-His tag and a TEV cleavage site and Cdc37 (residues 1-378) without tag were subcloned as the above-mention methods. The Flag-tagged Hsp90, the His_6_-tagged Cdk4 and Cdc37 was co-expressed in Sf9 cells using BAC-to-BAC Baculovirus Expression System (Thermo Fisher Scientific, USA).

#### Protein purification from Sf9 cells

The infected Sf9 cells were lysed and sonicated in 20 mM Tris-HCl (pH 7.5), 150 mM NaCl, 20 mM imidazole, 10 mM MgCl_2_, 10 mM KCl, 20 mM Na_2_MoO_4_. The protein solution was applied to a HisTap column (GE Healthcare, UK) and then eluted with a buffer containing 500 mM imidazole. Elution solution was applied to Anti-FLAG M2 agarose (SIGMA) and eluted a buffer (20 mM Tris-HCl (pH 7.5), 150 mM NaCl, 10 mM MgCl_2_, 10 mM KCl) with a 100 mg/ml FLAG peptide (SIGMA) to remove Cdc37 and Cdk4 unbound Hsp90. The Flag and His_6_ tag of the eluted sample was cleaved by TEV protease at 4°C for overnight. In turn, the solution was flowed through a HisTap column again to remove a cleaved Flag and His_6_ tag. The flow-through fraction was separated by an ion exchange column (MonoQ, GE Healthcare) and a size exclusion chromatography column (Superdex200, GE Healthcare) in a final buffer containing 20mM Tris-HCl (pH 7.5), 150 mM NaCl, 10 mM KCl, 10 mM MgCl2, 20 mM Na_2_MoO_4_, 2mM DTT. The sample was concentrated to final concentration of about 10 mg/ml and stored at −80°C until use.

#### Expression of the complex in *Saccharomyces cerevisiae*

To co-express human Hsp90β, human Cdc37 and human Cdk4 we utilized viral 2A peptides. This way we were able to construct a single plasmid, which had all three proteins in it. The exact 2A sequence we used (P2A) was sourced from Porcine Teschovirus-1 and was GSGATNFSLLKQAGDVEENPGP(*36*). The resulting construct was of this arrangement: hCdc37-TEVsite-P2A-hHsp90β-TEVsite-FLAG-P2A-hCdk4-TEVsite-HisTag. This construct was generated and cloned into 83nu yeast expression vector (GAL1 promoter, His marker) using Gibson Assembly(NEB). For T4 Lysozyme tagging T4Lys was added at either N terminus of Cdc37 or C terminus of Cdk4, separated from proteins by Gly-Ser (cloning was done at 96Proteins, South San Francisco). The resulting plasmids were sequence verified, transformed into JEL1 (MAT-alpha, leu2 trp1 ura3-52 prb1-1122 pep4-3 deltahis3∷PGAL10-GAL4) yeast strain using Zymo Research EZ Transformation protocol and plated on SD-HIS plates. After 3 days a colony was picked and 250mL O/N culture (SD-His) was inoculated. Next day 1L of YPGL media was inoculated with 10mLs of the O/N culture. After about 24h (OD of 1), galactose was added to a final concentration of 2% w/v to induce protein expression. The culture was pelleted after 6h of growth.

#### Purification of the complex from *Saccharomyces cerevisiae*

The cells were lysed in 20mM Tris pH7.5, 150mM NaCl, 20mM Imidazole, 10mM MgCl_2_, 10mM KCl, 20mM NaMoO_4_ (Lysis Buffer) with Roche protease inhibitors by Emusiflex (Avestin). Lysate was cleared by centrifuging at 30000g for 30 minutes and bound to pre-equilibrated Ni-NTA beads (ThermoFisher) for 1h at 4°C. The beads were washed with 20 bed volumes of Lysis buffer, and then were eluted into Lysis buffer + 500mM Imidazole. The resulting eluate was then incubated with pre-equilibrated M2 Anti FLAG magnetic beads (Sigma Aldrich) for 1h at 4°C. Beads were washed with 10 bed volumes of Lysis buffer, and the sample was eluted with 3 bed volumes of Lysis buffer with 75ug/mL of FLAG peptide, twice. TEV was added to the eluent and it was dialized against 20mM Tris pH7.5, 100mM NaCl, 10mM MgCl_2_, 10mM KCl, 20mM NaMoO_4_ (Dialysis Buffer) O/N. The sample was then diluted 1:1 with the Dialysis buffer without NaCl and was loaded onto pre-equilibrated 10/300 MonoQ column (GE Healthcare). After washing out the unbound sample, a gradient was run up to 1M NaCl with fractionation to elute the bound complex (came off at about 25% conductivity). The fractions were pooled, concentrated and then loaded on 16/60 S200 Superdex (GE Healthcare) column pre-equilibrated in 20mM Tris pH7.5, 150mM NaCl, 10mM KCl, 20mM NaMoO_4_, 1mM DTT. The peak fractions (at about 0.5CV) were pooled, concentrated, flash frozen in liquid nitrogen and stored at −80°C.

### Work on Cdc37-NTD expressed in *E.coli*

#### Expression and purification of Cdc37-NTD from *E.coli*

DNA corresponding to residues 1-126 of human Cdc37 was codon optimized for bacterial expression and ordered from ThermoFisher. Subsequently this construct was cloned into pet28a vector via Gibson assembly. Bl21Star (DE3) cells were transformed and plated. Colony was picked and an overnight culture in LB with antibiotic was grown. 6L of LB media with antibiotic were spiked with the O/N culture and induced at OD ~0.6 with 400uM of IPTG. After 3h of growth with shaking at 37C cells were pelleted by centrifugation. Cell pellets were solubilized in Lysis buffer (50mM Tris pH7.5, 500mM NaCl, 20mM Imidazole, 5mM BMe) and lysed on Emulciflex. The lysate was clarified by centrifuging at 30000g for 30 minutes, and the supernatant was collected. The supernatant was incubated with pre-equilibrated in Lysis buffer Ni-NTA beads (ThermoFisher) for 1h at 4C, rotating. The sample was eluted of the beads with Lysis buffer with 500mM Imidazole. TEV was added and the eluent was dialyzed against Dialysis Buffer (20mM Tris pH8, 10mM NaCl, 1mM DTT) at 4C O/N. The sample was diluted 1:1 with Dialysis buffer without salt and loaded on 10/300 MonoQ column. Protein was eluted by running a gradient to 1M NaCl over 20 column volumes. Appropriate peaks were collected and subsequently loaded on 16/60 Superdex 200 column. Appropriate peaks were collected, concentrated and flash frozen. For N^15^ growth the protocol was the same, except M9 minimal media with 1g/L N^15^ ammonium sulfate was used instead of LB during growth and expression.

#### NMR measurement on Cdc37-NTD

HSQC measurements were performed on a Bruker Avance 800 on fully N^15^ labeled Cdc37-NTD. Chemical shifts were visualized in ccpNMR.

#### CD measurement on Cdc37-NTD

Cdc37-NTD was buffer exchanged into 10mM Potassium Phosphate pH7.4, 50mM sodium sulfate buffer. Protein was diluted to 0.4 mg/ml and 1mm CD cuvette was used. CD spectra were collected on Jasco J710 from 185nm to 260nm.

### Cryo-EM data acquisition

#### Main data collection

Initially all the samples were screened using negative stain via standard protocols (~100nM protein concentration)(*37*). Cryo-EM grids were prepared with Vitrobot Mark III (FEI Company), using 20°C and 90% humidity. 3uL aliquots of sample at concentration of 1.1uM were applied to glow discharged C-flat 400 mesh 1.2/1.3 thick carbon grids (Protochips), single blotted for 4 to 6 sec and plunge frozen in liquid ethane cooled by liquid nitrogen. DDM was added to the protein to a final concentration of 0.085mM before applying sample to the grids. The grids with T4 Lysozyme labeled complex were prepared the same as above. Images were taken at NRAMM Scripps on FEI Titan Krios electron microscope operating at 300kV with a nominal magnification of 22500x. Images were recorded by Gatan K2 Summit detector (Gatan Company) with super resolution mode(0.66Å/pix). Defocus varied from 1.4um to 3.8um. Each image was fractionated to 38 frames (0.2sec each, total exposure of 7.6 seconds) with dose rate of 5.8e/Å/sec for a total dose of 44e/Å^2^. Leginon software was used for all the data collection. 3718 total images were collected. For more details see Table S1.

#### T4 Lysozyme labeled complex data collection

FEI Polara microscope operating at 300kV, at nominal magnification of 31000x, was used for data collection. Images were recorded with Gatan K2 Summit detector in Super Resolution mode (0.61Å/pix). Each image was fractionated into 30 frames (0.2 seconds each, 6 seconds total) at 6.7/Å^2^/sec for a total dose of 40e/Å^2^. Leginon software was used for all the data collection. For more details see Table S1.

### Image processing

#### Generating an initial reconstruction

Using EMAN2(*38*) for all aspects of processing, including initial model generation (e2initialmodel), a 3D reconstruction was generated from about 10000 negative stain particles. As mentioned in the main text, over multiple cryoEM data collections on F30 Polara, a 7Å reconstruction was obtained, using negative stain map as an initial model. The collection parameters were the same as T4 Lysozyme collections, but using UCSFImage4. This reconstruction, low pass filtered to 30Å, was used for particle picking and as an initial model for the NRAMM reconstruction.

#### General processing and obtaining the 3.9Å and 4Å reconstructions

Image stacks were corrected for motion and summed as described previously(*39*), resulting in binned sums (1.315Å/pix). For particle picking the images were binned to 5.2Å/pix and Gaussian bandpass filtered between 15Å and 500Å using EMAN2. SamViewer template based picking was then used to pick particles from all the micrographs, followed by manual review of all the picks(*40*). After such procedure 802877 particles were picked in total and extracted from images binned to 2.6Å/pix. CTFFIND4 was used to estimate defocus parameters for all the images(*41*). Relion 1.4 was used for all the following steps unless noted otherwise(*42*). Reference free 2D classification into 300 classes for 75 iterations was performed followed by manual examination of the resulting class averages. Low resolution/signal to noise/feature class averages and contributing particles were discarded, resulting in 670000 particles left. The resulting particles were 3D classified into 4 classes resulting in two classes having high-resolution features (390000 particles). At this stage particles were extracted from 1.315Å/pix micrographs and all the following processing was done with these particles. Using 3D Auto-refine in Relion 1.4, a reconstruction was obtained from 390000 particles resulting from 3D classification above (using highest resolution 3D class as initial model, low pass filtered to 20Å). Using the resulting parameters, the particles were further drift corrected per particle and dose weighted using the Particle Polishing feature(*43*). The B-factor weighing curve was fit by a polynomial (with a rationale that such a curve should be smooth) and used to generate new weighting parameters for Particle Polishing, with which 390000 particles were then polished. All further data processing was done using the polished particles. Re-refinement of the 390000 particles after polishing yielded the map at about 4Å resolution (determined using gold standard FSC in the PostProcessing tab)(*44*). Raw particles were sharpened with a B-factor of −50, low pass filtered with Gaussian filter to 3Å and the refinement was continued for 10 more iterations (until convergence) with these particles (the rationale was that due to extremely low noise levels of K2 direct detector, this would yield more accurate alignments due to presence of more high resolution data in the images). This resulted in similar resolution but a reconstruction with visually sharper features. This reconstruction, after Modulation Transfer Function (MTF) correction and B-factor sharpening(*45*) yielded the 4Å reconstruction shown on Fig1 and shown without post processing in Fig2A and movie S2. Lastly, this reconstruction was tightly masked around Hsp90 region and refinement was further continued for 4 iterations (until convergence), with rationale that Hsp90 region is more coherent. This yielded a 3.9Å reconstruction with the best density for Hsp90 region (Shown in inserts on Fig 1B, 2B and Fig 5), at the expense of Cdk4 C-lobe and Cdc37 NTD density quality. Local resolution was estimated using ResMap(*46*).

#### Obtaining 3D classes for Cdc37 and Cdk4 N-lobe

The 390000 polished particles were further sub-classified into four 3D classes. All four resulting reconstructions were better than at 10Å resolution, with Hsp90 density staying unchanged, but the densities on the periphery changing. One of the classes had a low-resolution density where Cdc37 M/C was placed later, but at this resolution the fit would be ambiguous. To obtain higher resolution reconstruction of this region, a local mask was generated around this region. The 390000 particles were then 3D classified into four different classes without particle re-alignment, using the alignment parameters from 4Å reconstruction. Particles contributing to each of the four classes were grouped and a full 3D refinement with a spherical 200Å mask was performed with each of the four groups of particles using the same initial model, low pass filtered to 20Å. One of the classes (referred to as Cdc37 Reconstruction throughout the text and shown on Fig 3A) had a distinct density into which we fit Cdc37 M/C (in UCSF Chimera(*47*)). There was also a low-resolution density for the kinase N-lobe. To attain a better density for the kinase N-lobe, we re-did the local masked 3D classification as described for Cdc37 Reconstruction, but used the subtraction protocol described recently(*48*). Again, particles contributing to each of the masked classes were grouped and a full reconstruction with a spherical 200Å mask was performed for each of the groups, starting with identical initial models. Such a procedure generated the two kinase reconstructions (Blue Kinase reconstruction and Maroon Kinase reconstruction shown in Fig3 B), a reconstruction with Cdc37 M/C density (but no Cdk4 N-lobe density) and a reconstruction which had no extra densities in that region (similarly to the 4Å reconstruction). The last two reconstruction of this set were not show or used in this manuscript. All the reconstructions were filtered and sharpened using the PostProcessing tab in Relion. All the reconstructions were visualized using UCSF Chimera.

#### Processing of T4 Lysozyme tagged data

Image stacks were dose weighted, drift corrected, binned to 1.22Å/pix and summed using new UCSF DriftCorr program. The particles were picked and the CTF was estimated the same way as in the main data collection. Rounds of 2D classification (300 classes, 50 iterations) followed by 3D classification (2 classes, 50 iterations) in Relion 1.4 were used to eliminate low quality particles (using 30Å low pass filtered reconstruction from the main data set as initial model). Using the final set of particles, 3D Auto-refine feature in Relion 1.4 was used to generate the final maps. The parameters of final reconstructions and post processing are reported in Table S1.

### Model Building and refinement

Atomic model building and refinement the Hsp90/Cdc37/Cdk4 complex was performed incrementally in five stages: 1) *de novo* model-building for Cdc37, 2) structure refinement of the Hsp90/Cdc37 complex, 3) *de novo* model extension for Cdk4 in the presence of the refined Hsp90/Cdc37 complex, and 4) structure refinement of the Hsp90/Cdc37/Cdk4 complex. The atomic structure of Hsp90/Cdc37/Cdk4 complex was used in the modeling of other low-resolution maps.

#### Initial fitting of hHsp90/Cdk4 C-lobe model

The V-shape structure of Hsp90 was clearly identified from the 4Å reconstruction, in which it adapts the closed-state conformation of known Hsp90 structure. The hHsp90 homology model was derived from a close (~60% sequence identity) homologous structure from *Saccharomyces cerevisiae* (PDB: 2CG9 chain A and B), which also adapts a closed-state conformation. The unrefined hHsp90β model was first rigid-body fit into the map using UCSF Chimera (Fit In Map function), revealing good agreement with the density data, where most of the secondary-structure density features can already be explained. The Cdk4 C-lobe (residues 96-295) from PDB:3G33 was fit based on structural alignment to a CATH(*49*) domain 3orkA02. This CATH domain was obtained by downloading all the CATH protein folds and running COLORES program (Situs package)(*50*) on each, rigid body fitting it into the segmented globular region density (Fig 1B arrows and Fig S6). This resulted in an un-biased fit into the density. The initial hHsp90β homology model was built together with Cdk4 C-lobe using the Molecular Dynamics Flexible Fitting (MDFF) (*51*).

#### *De novo* building Cdc37-NTD into 4Å reconstruction using Rosetta

With no homologues of known structure available for the Cdc37 N-terminal domain (Cdc37-NTD), we employed the Rosetta automated *de novo* model-building method (*52*) to register the sequence (residues 1-132) in the density. The density belonging to Cdc37-NTD was manually segmented guided by the initial hHsp90/Cdk4 C-lobe model using Chimera’s “Volume Cleaner” tool. The method first places 9-mer fragments derived from local sequence into the density map, in order to simultaneously trace the backbone and assign sequence. High-confidence partial models are generated using Monte Carlo sampling to identify a set of placed fragments consistent with each other and with maximum agreement to the data. After two iterations of applying the procedure described above, the method converged on a partial model with residues ranging from 2-50 and 92-122. RosettaCM was used to complete the partial model through density-guided model rebuilding and refinement (*53*). To this end, *de novo* model-building of Cdc37-NTD was done in the segmented density alone. However, it was clear from our initial model that Cdc37 had extensive contacts with hHsp90 and Cdk4 C-lobe in the map. Therefore, we carried out a further structure refinement in the context of Cdc37-NTD and hHsp90.

#### Refinement of Hsp90/Cdc37 into 3.9Å map using Rosetta

The starting model of hHsp90 and Cdc37-NTD was obtained as described in the previous paragraph. A split map approach was used to prevent and monitor models from data over-fitting, and for model selection; one of the half maps used for calculating “gold-standard” FSC was designated as the training map against which the model was refined, and the other half map was designated as the testing map which was used for evaluations. The training/testing map designations were strictly followed and consistent in all stages of refinement. Guided by the starting Hsp90/Cdc37 model, density from the Cdk4 and nucleotide was carefully removed from the training map using Chimera’s “Volume Cleaner” tool.

The refinement procedure has five cycles of iterative backbone rebuilding; in each cycle, regions with poor fit to density or poor local geometry were automatically identified, and rebuilding focused on these regions. Backbone rebuilding of a residue identified as problematic is similar to what has been described in DiMaio et al.(*54*), where the rebuilding starts with replacing the coordinates of the current model with fragments at the residue position through Monte Carlo sampling, followed by Cartesian space minimization. The fragment with the best fit to the local density is selected; the backbone coordinates from the current model are replaced by the selected fragment. Each rebuilding cycle was followed by side-chain rotamer optimization and all-atom refinement with a physically realistic force field to ensure the regularity of the rebuilt backbone. After the backbone rebuilding cycles, a “dual-space” all-atom refinement which alternates between torsion-space and Cartesian-space optimization was carried out, and was followed by a newly-developed “local relax” to ensure the convergence of side-chain rotamer optimization in a large protein complex. Following this procedure, ~5,000 independent trajectories (models) were run.

Model selection was done in three steps. First, the initial pool of models were picked by selecting 30% most physically realistic models accessed by the Rosetta all-atom force field, from which the top 20% most well fit-to-density models were selected using Rosetta all-atom electron density score (“elec_fast_dens”). Second, evaluating the fit-to-density of the initial pool of models against the testing map using the high-resolution shells (10-3 Å) of FSC; 50 models with low over-fitting (FSC_training_ - FSC_testing_ < 3%) and high correlation with the testing map were retained. Third, 10 out of 50 models were picked using Molprobity (*55*), sorted first by Molprobity score and then “Ramachandran favored” score. Lastly, a final model was selected among the 10 models through visual inspection in Chimera. Here, the above described model refinement/selection procedure was carried out with two iterations to reach a satisfactory model. Residues with poor fit to density were specified to correct for the next iteration refinement.

#### *De novo* model extension for the docked Cdk4 C-lobe (into 3.9Å map)

Starting with a docked configuration of the Cdk4 C-lobe (residues 96-295), *de novo* building of the upstream residues 86-95 (tube) was carried out using RosettaCM into the tubular density as described in the main text. This density has extensive contacts with the other two component proteins, Hsp90 and Cdc37-NTD. Thus, model building into the tube density was done in the presence of Hsp90 and Cdc37-NTD. Density-guided conformation sampling was primarily focused on the tube and the residues of Hsp90 that were likely in contact with (residues 341-349 Hsp90 chain A and B), which combines Monte Carlo sampling of backbone fragments with Cartesian space minimization in the context of residues around the tube-like density. 50 low-energy models were selected by finding models with physically realistic energy using Rosetta all-atom energy, as well as good density-fit. Among the 50 low-energy models, one conformation was clearly favored and showed good agreement with the density. Next, we used the iterative backbone rebuilding procedure as described in the previous paragraph to further sample and optimize the tube residues, as well as the surrounding residues from Hsp90. Finally, the conformation again was favored and converged among the low-energy models.

#### Refinement of the Hsp90/Cdc37/Cdk4 complex in 3.9Å map

Refinement of the Hsp90/Cdc37/Cdk4 complex was done out using the same procedure described in the paragraph of refining the Hsp90/Cdc37 complex. However, due to the much worse resolution (~ 6 Å) of the density for Cdk4’s C-terminal domain (residues 96-295, chain K) and Cdc37-NTD residues 56-91 (chain E), it was not necessary to further optimize the structure; thus, no fragment-based rebuilding was allowed in these residues. Three iterations of automatic model refinement/selection procedure were carried out. Furthermore, through visual inspection residues that showed poor density-fit were pinpointed and subject to the iterative backbone rebuilding on only those residues. With three more iterations of human-guided model refinement/selection, a satisfactory model was finally obtained. Next, heteroatoms including nucleotides (ATP), Magnesium and phosphorylated-Serine were appended into the model. The intial ATP conformations were adapted from the yHsp90 crystal structure (PDBID: 2CG9). The model was refined using the Rosetta *relax* protocol with Cartesian space optimization, followed by the *local_relax*. The same model selection procedure described in the previous paragraph was used to select the final model of Hsp90/Cdc37/Cdk4 with heteroatoms. B-factors of each atom in the atomic model were refined in the full reconstruction using the approach from DiMaio et al.(*54*). Finally, we used the half maps to find a weight of density map that does not introduce over fitting. Using this weight, we did a *local_relax* in the 3.9Å map reconstructed from all the 390000 particles. The resulting model (hHsp90β 10-690, hCdc37 2-132, hCdk4 86-295) would be used as rigid body models for fitting into all the low resolution maps downstream.

#### Generating a model for residues 1-260 of Cdc37

Cdc37 crystal structure from PDB:1US7 was fit into the Cdc37 Reconstruction (not post processed, so, low pass filtered to ~7Å and no B-Factor sharpened) manually and then with UCSF Chimera “Fit In Map” tool. The model was truncated at residue 260, as there was no reliable density for the rest of the crystal structure. The Rosetta built model generated in the previous paragraph and the above fit crystal structure for residues 148-260 were loaded in Coot(*56*). The residues 133-147 were built in by hand into the Cdc37 Reconstruction and residues 245-260 (helix) were rotated as a rigid body. To relieve atomic clashes or bond length/angle distortions at the linker regions, the resulting model was subjected to “Cartesian space relax” protocol within Cdc37 Reconstruction density map using Rosetta. Final model was selected using the combined score of Rosetta all-atom physically-realistic score and electron density score.

#### Generating a complete Cdk4 model (Blue and Maroon Kinase reconstructions)

Residues 5-85 (N-lobe) of Cdk4 were fit in Chimera as rigid body into the appropriate density in each of the reconstructions. Such fit N-lobe of Cdk4 was further tweaked in Coot in the context of model generated in the previous paragraph to join the Cdk4 chain and minimize clashes for each of the maps. To relieve atomic clashes or bond length/angle distortions at the linker regions, these models were subjected to “Cartesian space relax” protocol within corresponding density maps using Rosetta. Final models were selected using the combined score of Rosetta all-atom physically-realistic score and electron density score.

#### Fitting of T4 Lysozyme into the T4 Lysozyme reconstructions

Hsp90/Cdc37/Cdk4 model was fit into each of reconstructions in UCSF Chimera. T4 Lysozyme (PDB:2LZM) was fit into the extra densities first by hand and then in UCSF Chimera. No further refinement was undertaken.

### PDB and EMDB accession numbers

The accession numbers are EMD-3337, EMD-3338, EMD-3339, EMD-3340, EMD-3341, EMD-3342, EMD-3343 and EMD-3344 for EMDB and 5FWK, 5FWP, 5FWL and 5FWM for PDB.

### Figure preparation

All model and reconstruction figures were prepared in UCSF Chimera except Fig. 4B which was prepared with pre-release version of ChimeraX.

## Acknowledgments

We thank members of NRAMM-Scripps for help collecting data, Yao Fan for the yeast expression vector, Naomi Ohbayashi and Mutsuko Niino (RIKEN Center for Life Science Technologies) for help with Sf9 protein expression, Dr. Neil F Rebbe (The University of North Carolina at Chapel Hill) and Dr. Ernest Laue (The University of Cambridge) for the plasmids encoding human HSP90b and Cdc37, respectively, D.A.A. lab members for helpful discussions and Natalia Jura for reading the manuscript. Support for this work was provided by PSI-Biology grant U01 GM098254 (to D.A.A.), AACR-BCRF Grant 218084 for Translational Breast Cancer Research (to D.A.A.), The Cabala Family gift (to D.A.A.), HHMI International Student Research Fellowship (to K.V.) and the Howard Hughes Medical Institute (to D.A.A.). Some of the work presented here was conducted at the National Resource for Automated Molecular Microscopy which is supported by a grant from the National Institute of General Medical Sciences (9 P41 GM103310) from the National Institutes of Health.

## Supplementary figures and materials

**Fig. S1.**
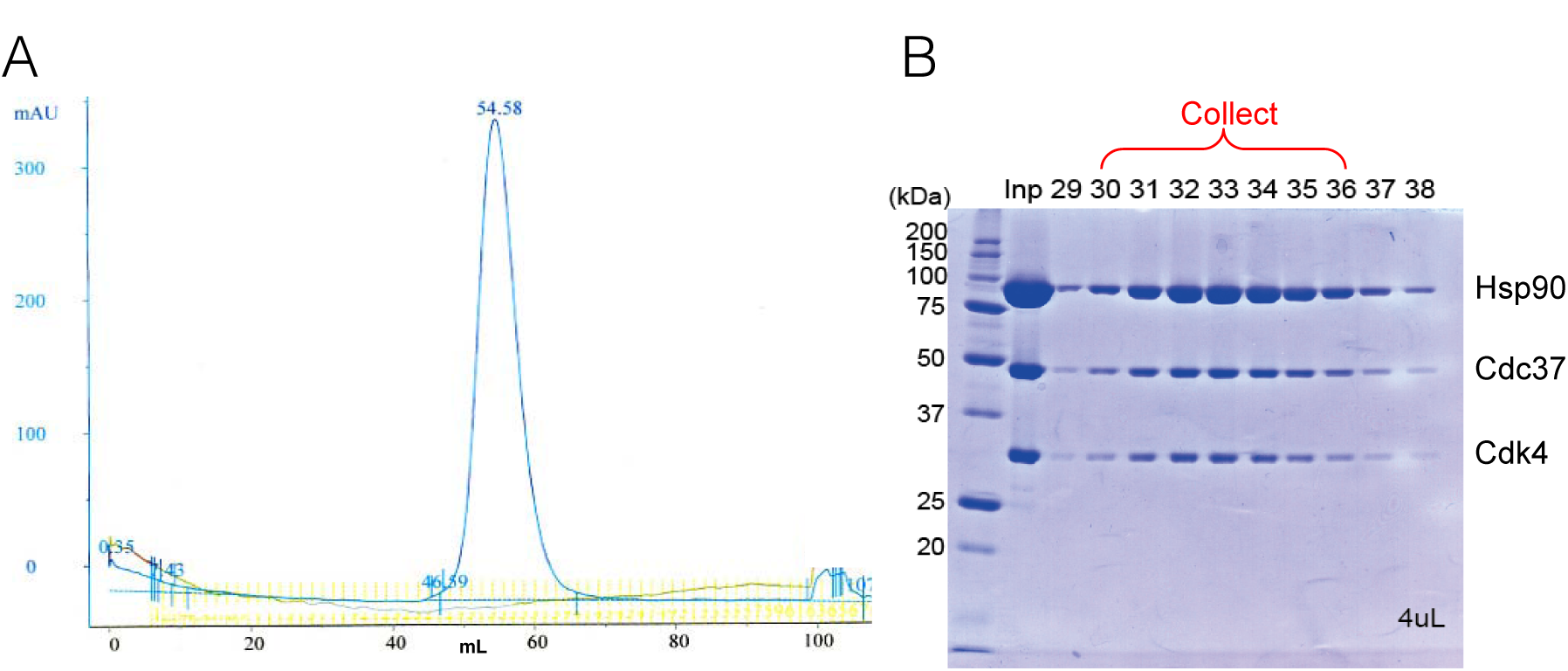
Purification of Hsp90/Cdc37/Cdk4 complex from Sf9 cells. (A) Final sample is monodisperse on S200 Superdex gel filtration column. (B) The peak from the gel filtration ran on SDS-PAGE gel showing that all proteins are present in the final sample arid are highly pure.

**Fig. S2.**
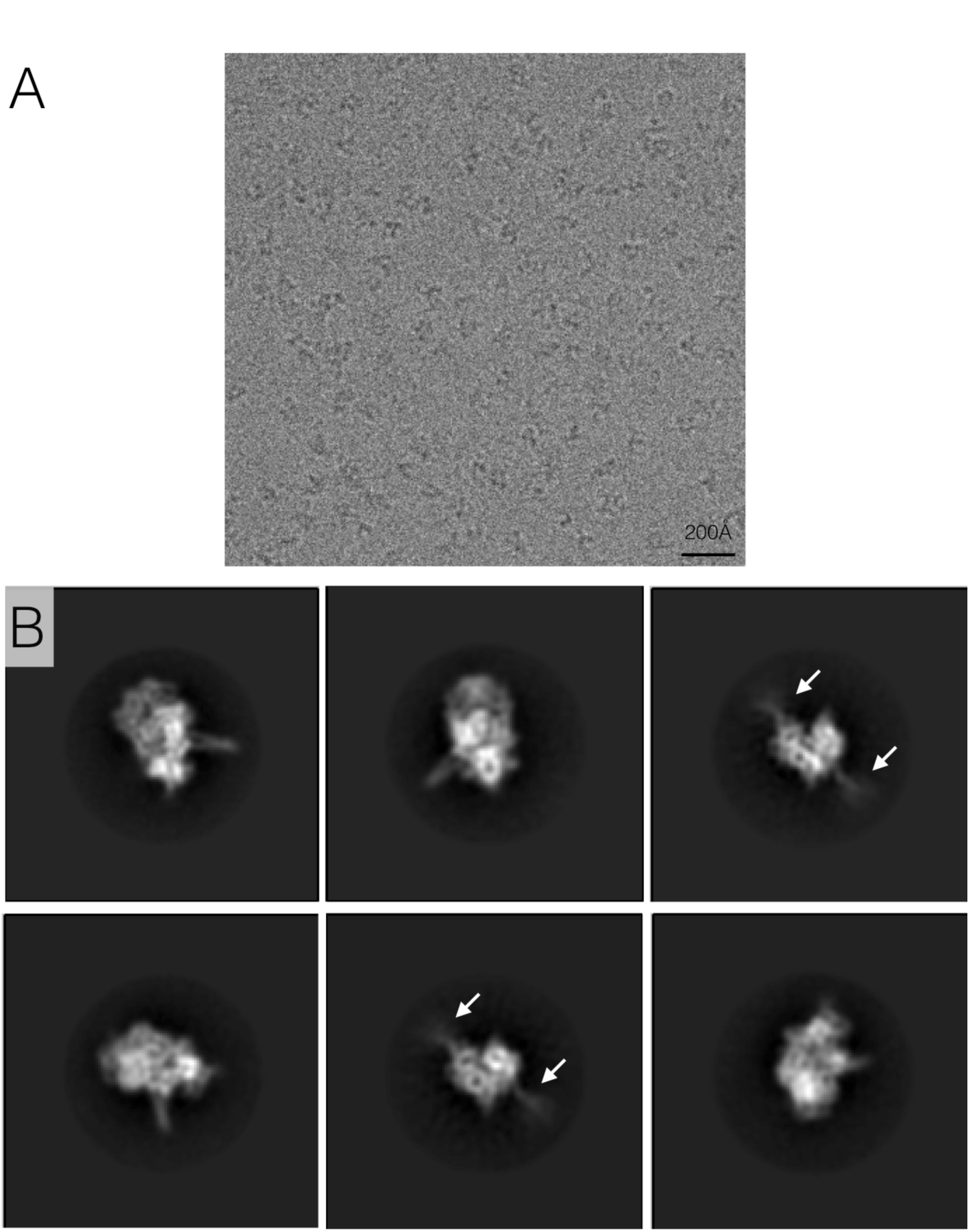
Pre reconstruction assessment shows high quality data. (A) A crop of unfiltered drift corrected image stack sum shows clearly visible particles. (B) 6 classes were picked from a total of 300 reference free classes (Relion). The classes clearly show secondary structure elements and the coiled coil protrusion. Marked with white arrows is the charged linker, which clearly becomes more diffuse the further it is from the center.

**Fig. S3.**
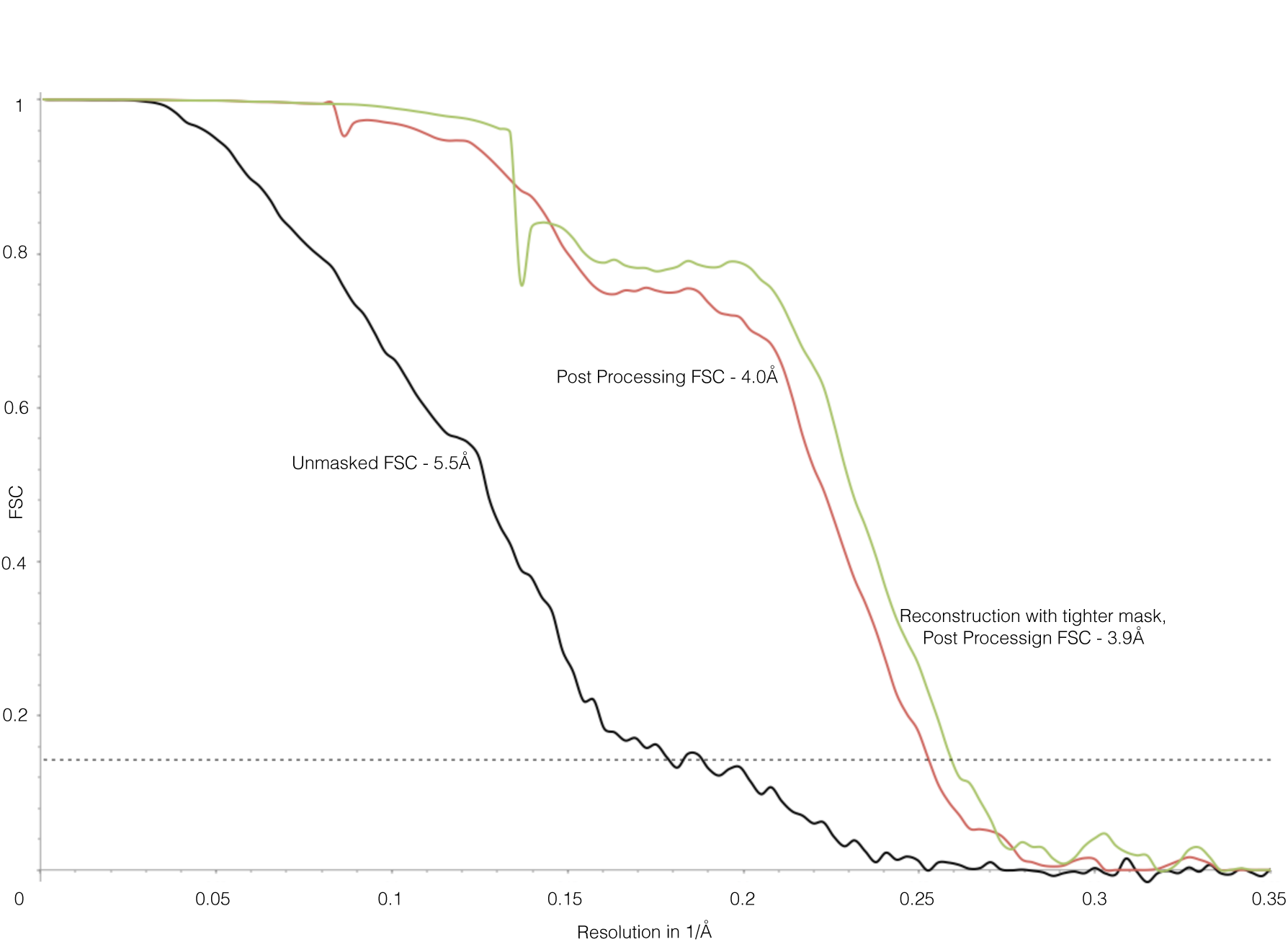
Gold standard FSC of our highest resolution reconstruction. In black is the unmasked FSC (ie 200Å spherical mask) of the reconstruction, crossing 0.143(dashed line) at 5.5Å. In red is the masked, corrected FSC of the same reconstruction, crossing the 0.143 threshold at 4.0Å, generated in the post processing node in Relion. In green is the corrected FSC for the 3.9Å reconstruction, which was performed utilizing a mask around Hsp90 region (See methods for more details).

**Fig. S4.**
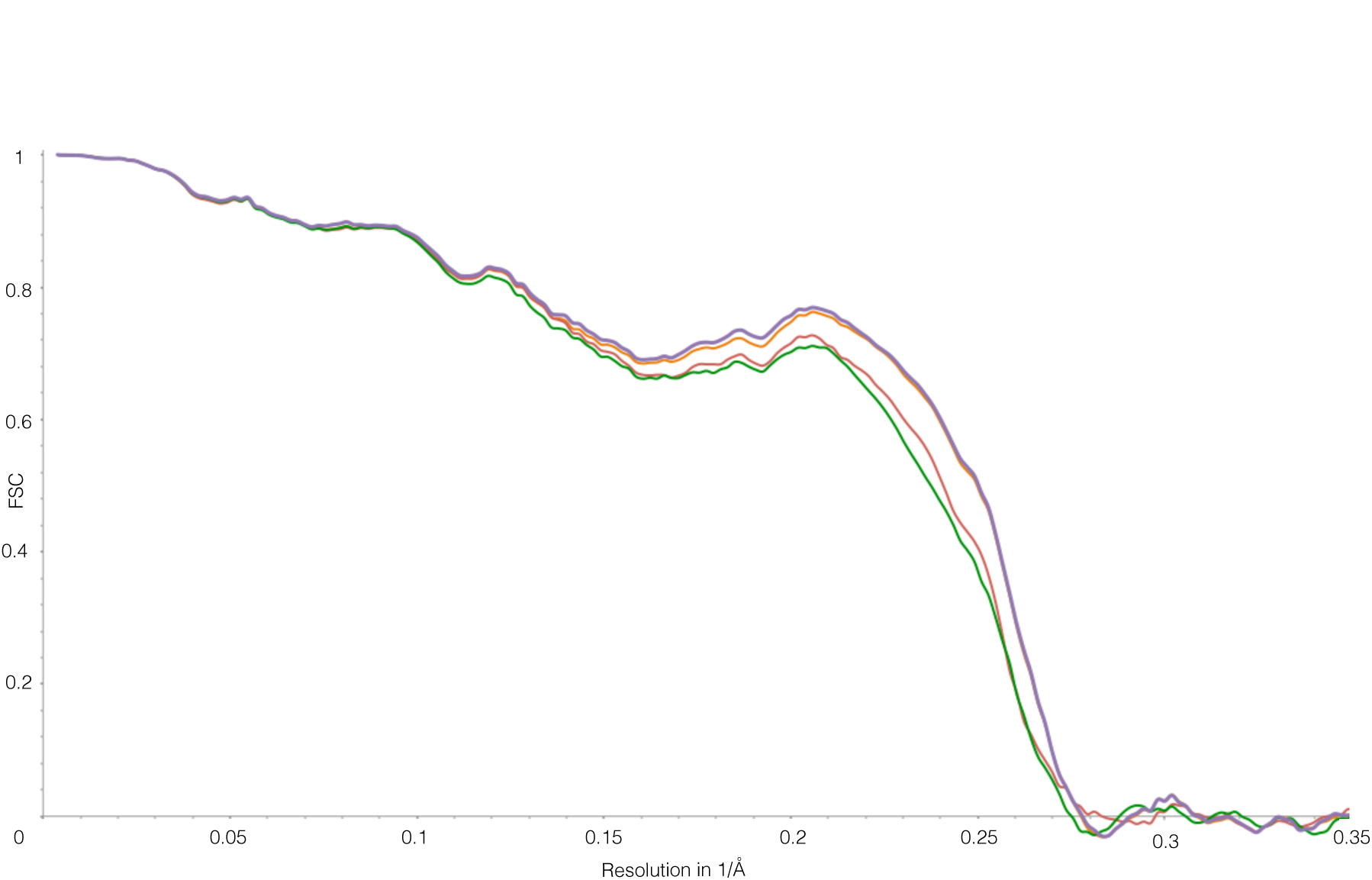
Model vs map FSC shows no over fitting. The difference between model vs the training map FSC (red curve) and model vs test map FSC (green curve) is small, signifying no over fitting. In orange is the FSC between the model refined into a half map and full 3.9Å map. In violet is the FSC of the final model vs 3.9Å map. The final map was refined into the map from full dataset of particles using weighting parameters from the half map refinements. (See methods for more details).

**Fig. S5.**
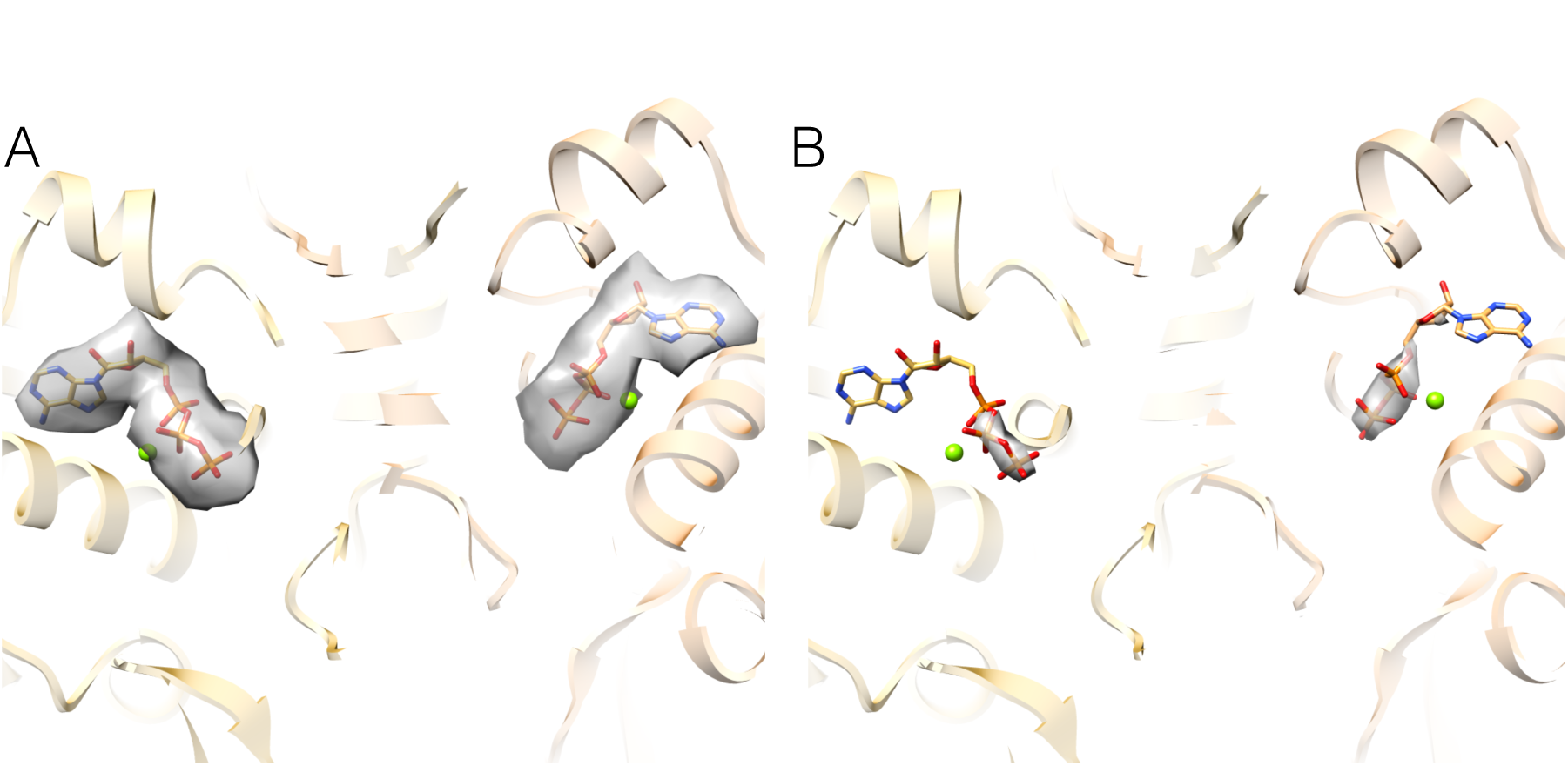
There is a clear density for nucleotide in both Hsp90 monomers. (A) View from the 4Å map showing the ATPs fitting perfectly into the density in both monomers. (B) The density for the γ-phosphatc is the strongest density in the whole map.

**Fig. S6.**
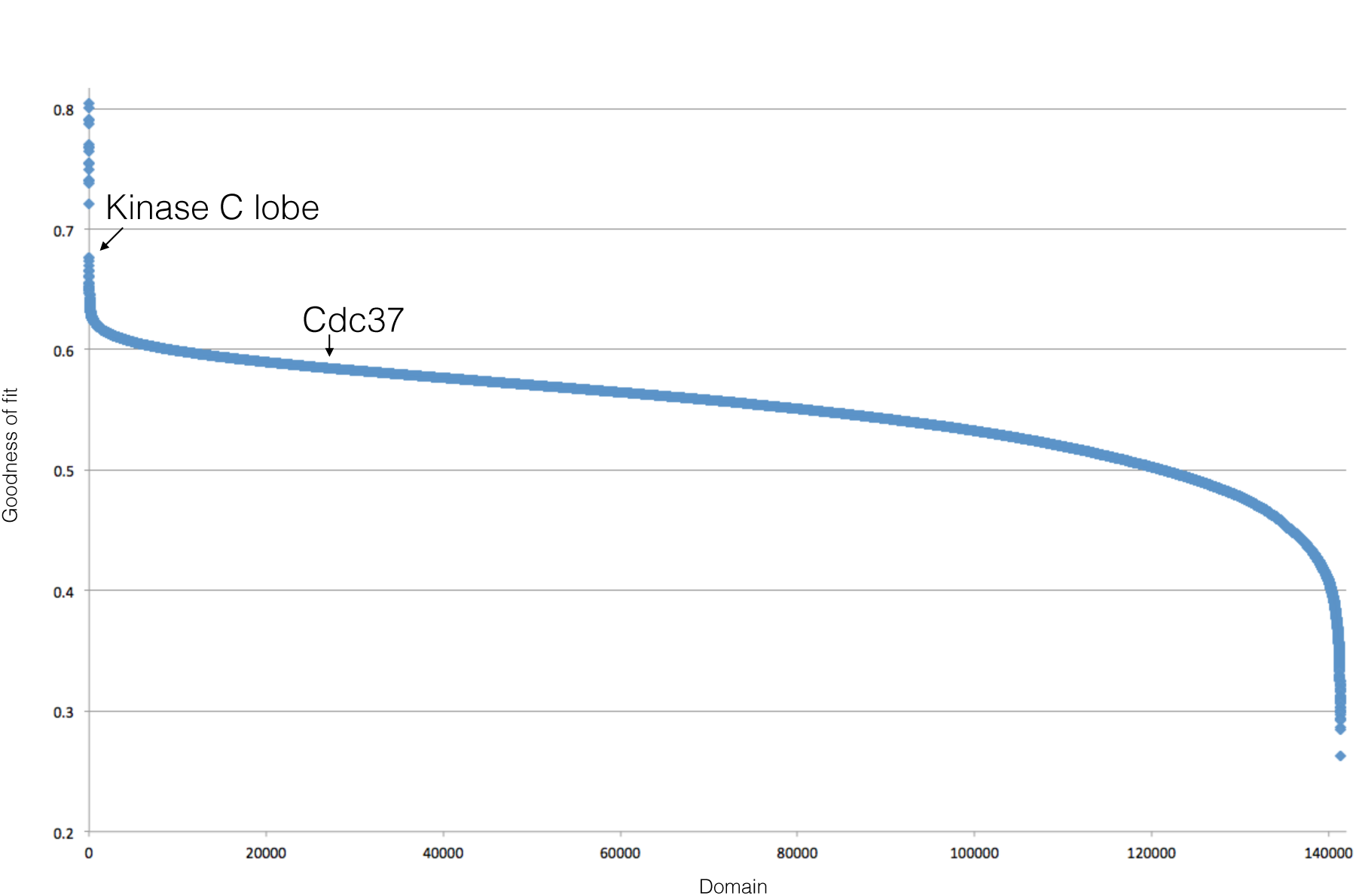
Fits of different CATH domains into the globular density plotted by cross correlation. The plot of the cross correlation coefficient vs CATH database domain, as output from COLORES program in SITUS package. Upon manual examination, all the points above 0.7CC are clearly artifacts. The CC values from kinase C lobe and Cdc37 are marked with arrows. There are more data points than CATH domains as COLORES outputs multiple alternate fits for each CATH domain.

**Fig. S7.**
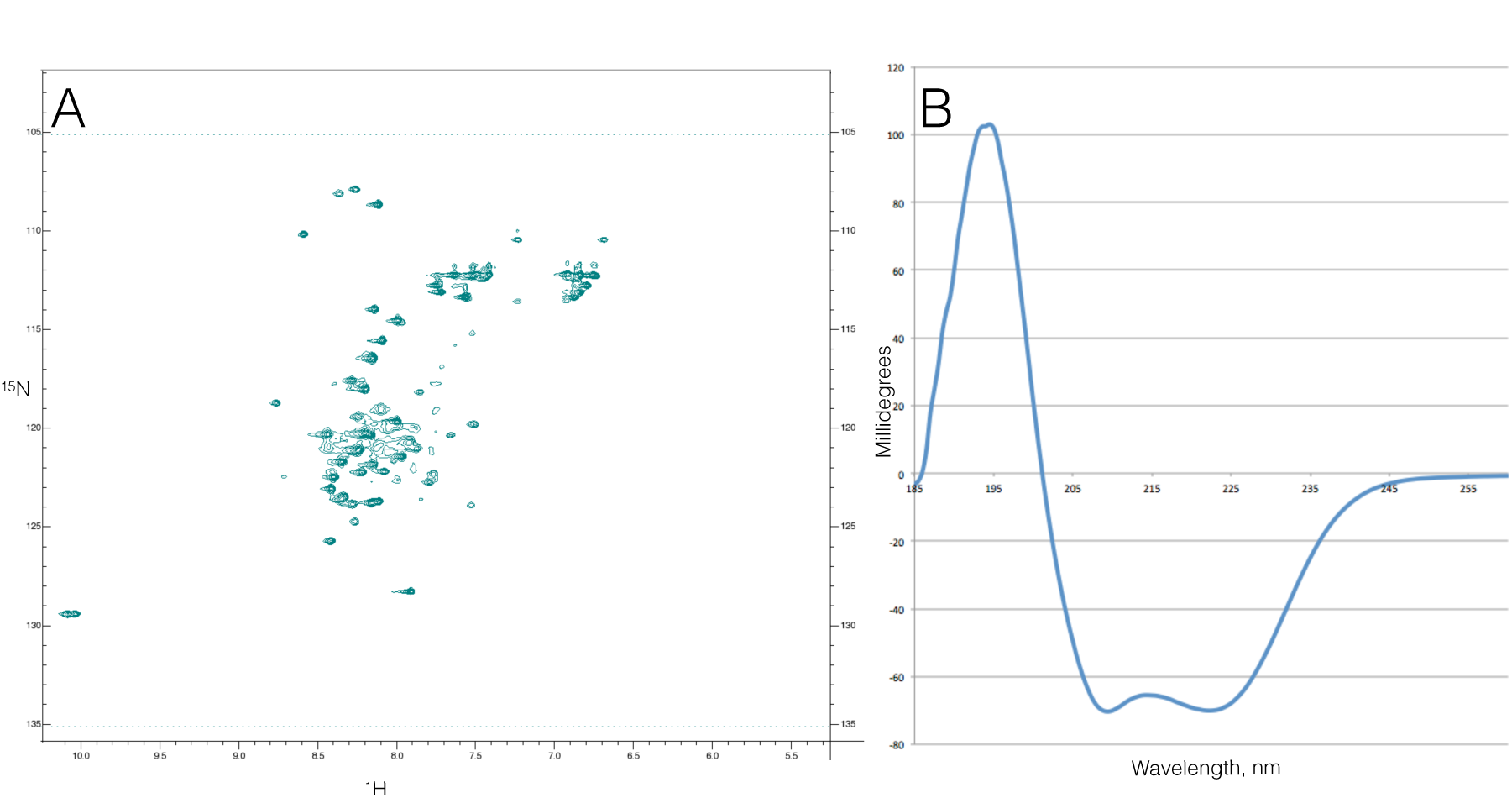
CDC37 N-terminal domain is helical. (A) NMR spectrum of N^15^-H^1^ HSQC experiment performed on the CDC37 NTD, which is characteristic for either unfolded or helical proteins. (B) CD of the same protein fragment, showing spectrum characteristic of α-helix.

**Fig. S8.**
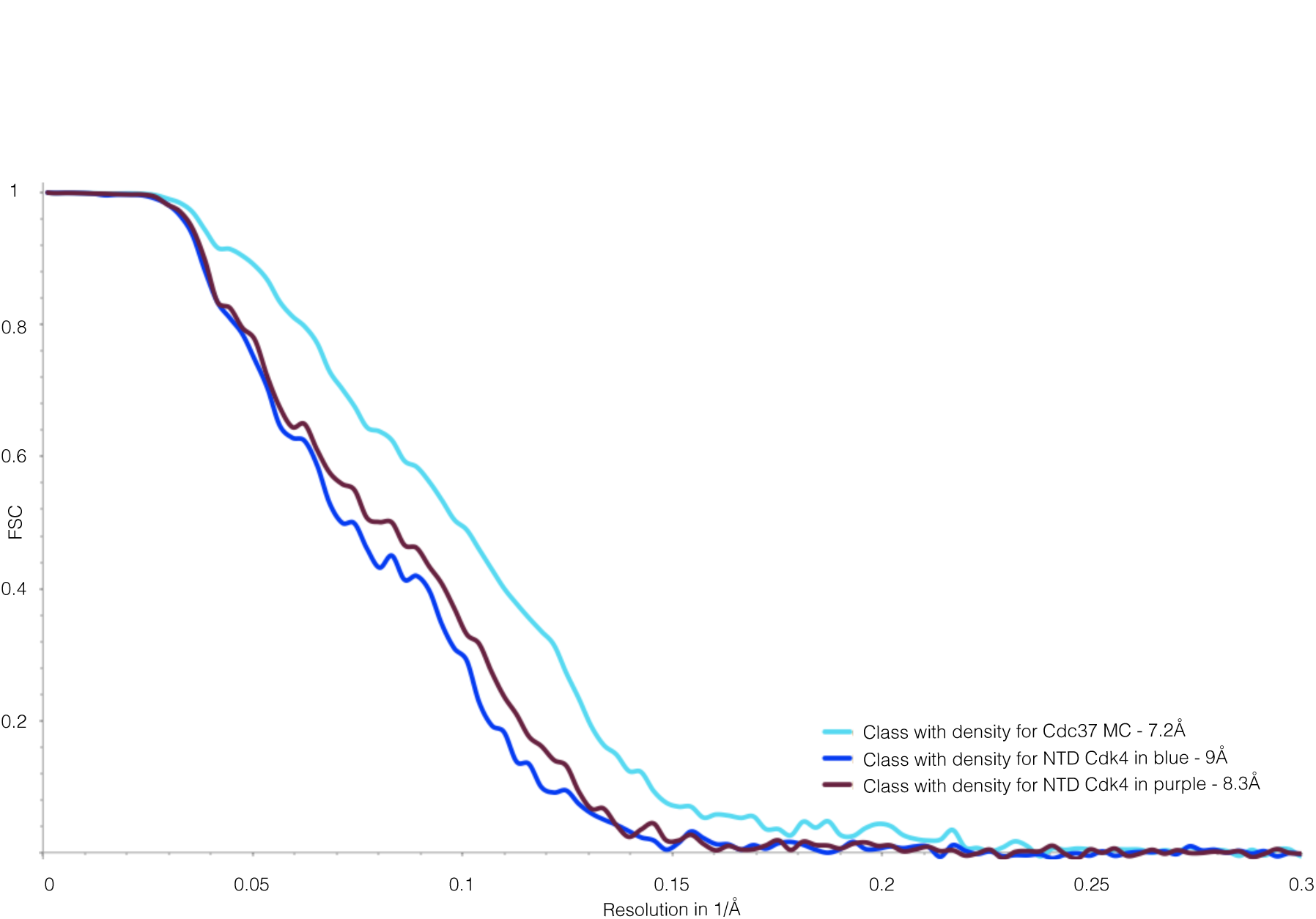
Gold standard FSCs of the three alternate reconstructions. Unmasked gold standard FSCs of the three reconstructions done after local 3D classifications, as discussed in the main text. Due to using only a spherical mask, the resolution estimates are quite conservative.

**Fig. S9.**
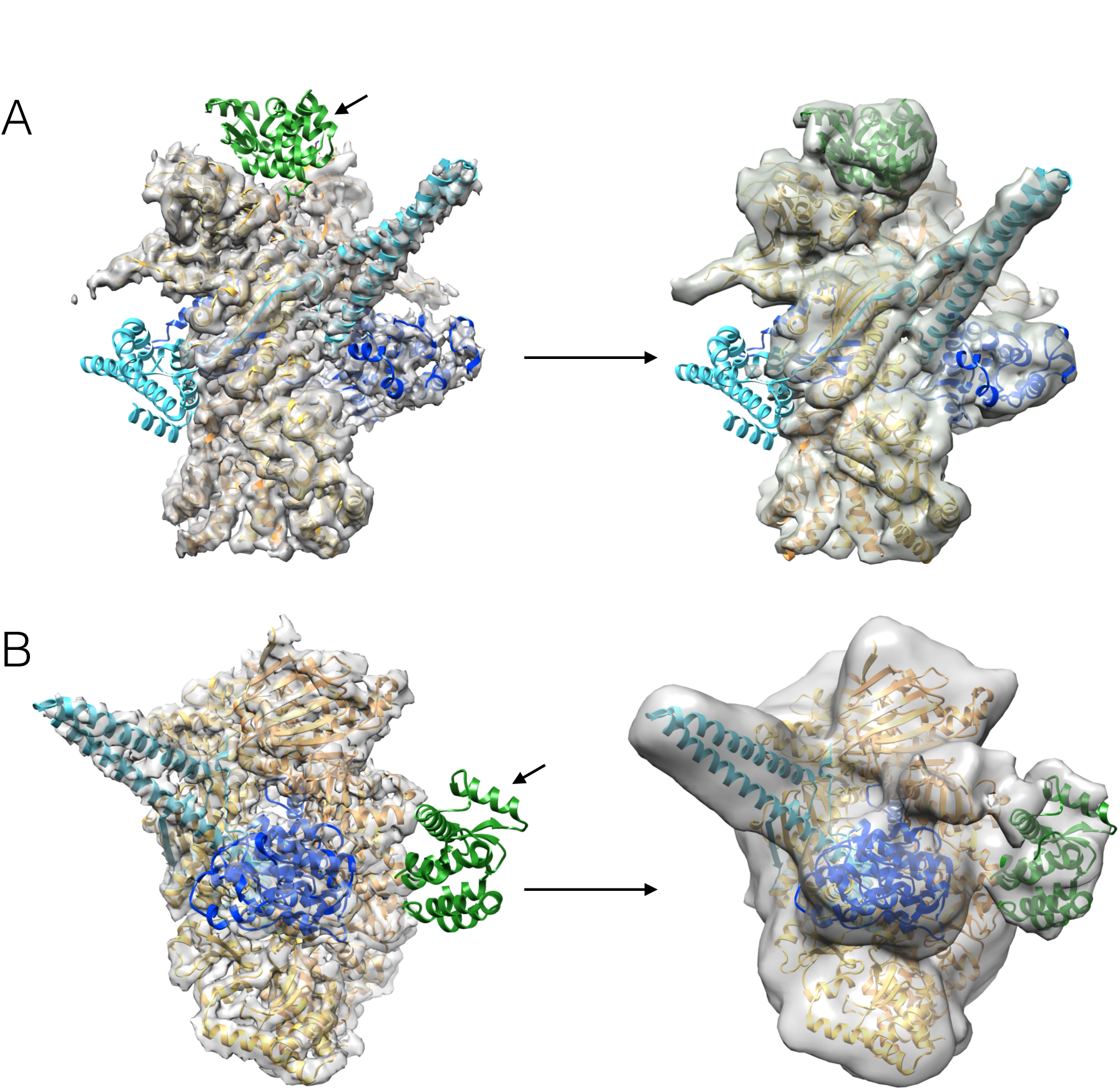
T4 lysozyme tagging supports protein placement in our model. (A) Cdc37 was tagged on its N terminus with T4 lysozyme. On the left is our model fit into the 4Å map, with lysozyme (PDB:2LZM) in green placed at its expected location as per construct(marked with an arrow). On the right is the reconstruction of the Hsp90/T4Lys-Cdc37/Cdk4 complex, clearly showing the new extra density in the expected location perfectly fitting lysozyme structure. (B) Everything as in A, but now Cdk4 was tagged with T4 Lysozyme at its C terminus. Again, reconstruction of Hsp90/Cdc37/Cdk4-T4Lys shows a clear new extra density for lysozyme.

**Fig. S10.**
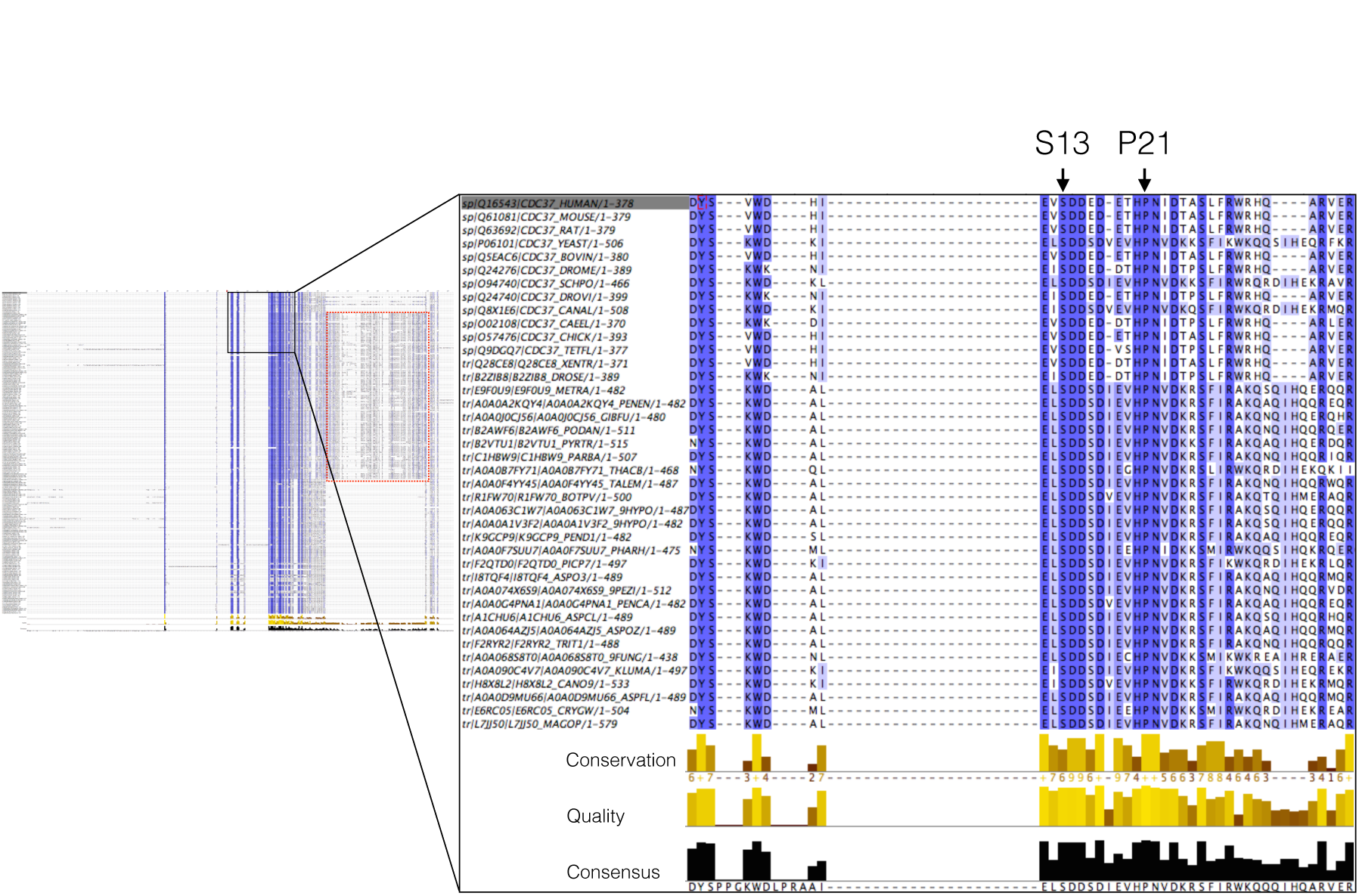
Cdc37s very N terminus is extremely well conserved. Sequence alignment of over 200 different Cdc37 protein sequences aligned using T-COFFHH server and visualized using JalView. Red rectangle is showing a long insertion characteristic of fungi Cdc37. The insert is showing a zoom view of the first 30 amino acids of 40 organisms, with the top sequence being human. Conservation in the region is over 90% for many of the residues.

**Fig. S11.**
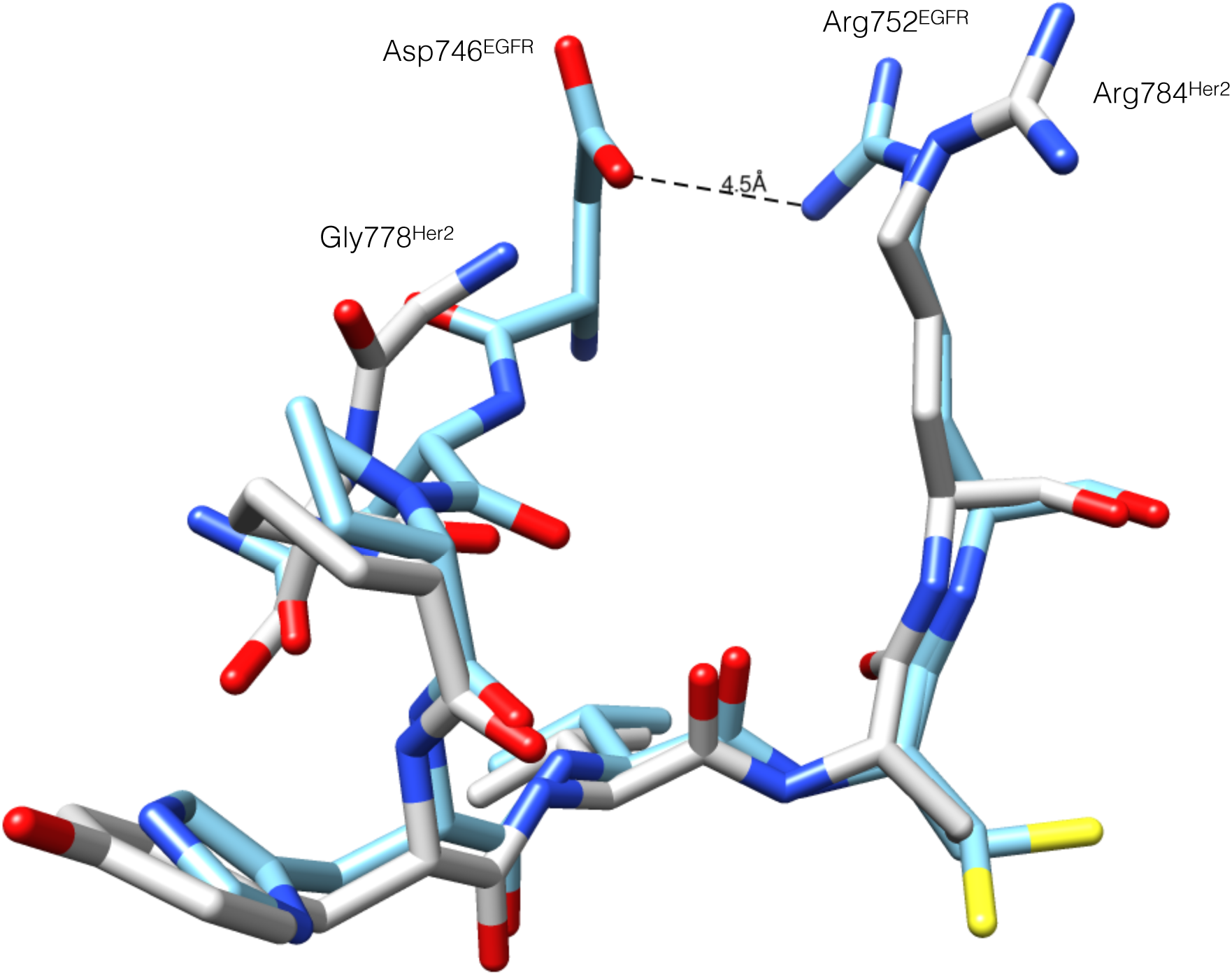
αC-β4 loop in EGFR may be stabilized by an ionic interaction. Overlay of the loop between Her2 and EGFR structures, with Her2 being in white and EGFR being in blue. Considering the overall flexibility of protein kinases, Asp746 in EGFR may electrostatically interact with Arg752, stabilizing the loop. This interaction would be absent in Her2 as Asp746 is replaced with Gly778. (PDB codes 3PP0 and 1M17)

**Table S1.**
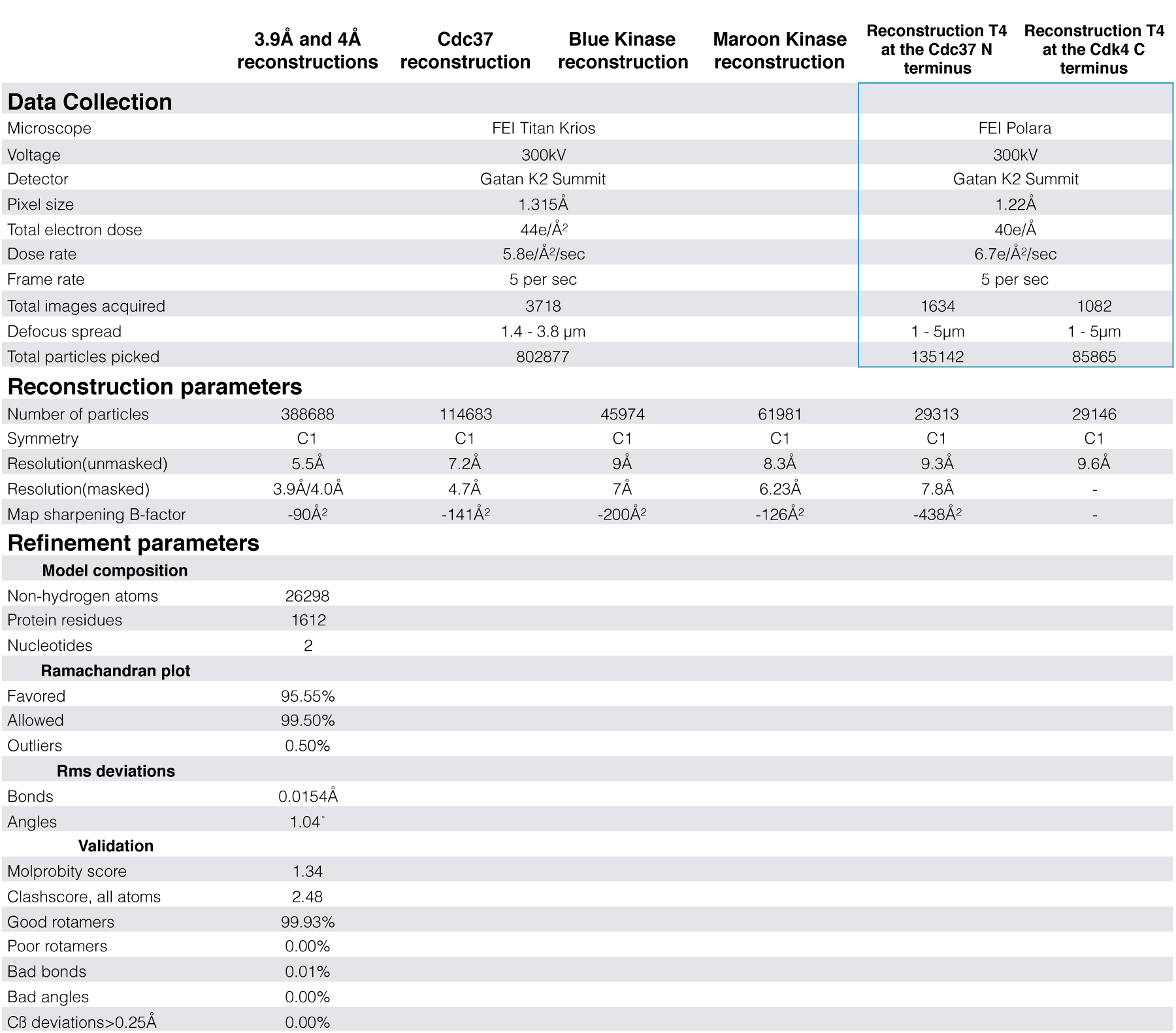
Parameters for all the reconstructions discussed in the main text and refinement parameters for the final model refinement (omitting Cdc37 M/C and Cdk4 N-lobe) fit into the 3.9Å reconstruction.

**Table S2.** Buried surface area between the proteins in the complex, per domain, generated by submitting the final model into the PISA server.

| Protein Fragment 1 | Protein Fragment 2 | Interface Area (Å<sup>2</sup>) |
| --- | --- | --- |
| Cdk4 | Hsp90 A | 639 |
|  | Hsp90 B | 922 |
|  | Hsp90 NTD A | 90 |
|  | Hsp90 MD A | 225 |
|  | Hsp90 MD B | 572 |
|  | Hsp90 CTD A | 366 |
|  | Hsp90 CTD B | 376 |
|  | Cdc37 | 1001 |
|  | Cdc37NTD | 725 |
|  | Cdc37MC | 300 |
| Cdc37 | Hsp90 A | 654 |
|  | Hsp90 B | 2000 |
| Cdc37NTD | Hsp90 NTD A | 230 |
|  | Hsp90 NTD B | 67 |
|  | Hsp90 MD B | 1608 |
| Cdc37MC | Hsp90 MD B | 337 |
| Cdc37MC | Hsp90 CTD A | 424 |

**Movie S1 Movie showing Hsp90s CTD rotation to accommodate client binding.** Hsp90 and Cdk4 are colored as before. The movie starts with 2CG9 like state and ends with the state in our structure. The regions that are client interacting and are ordered in our structure are in magenta. They are showed as disordered in the movie due to Morph Conformations methodology in Chimera.

**Movie S2 Movie focusing on the Cdk4 C-lobe and part of N-lobe going through the lumen of Hsp90.** In the light blue is a structure of a C-lobe from the CATH database, in medium blue is a structure of full length Cdk4 and in dark blue is the structure of Cdk4 with the N-lobe unfolded. The map which was used is the 4Å reconstruction without post processing (ie, no b-factor sharpening, filtered to 5.5Å)

**Movie S3 Movie showing morphs between 4 different reconstructions discussed in the text.** In gray is the 4Å reconstruction filtered to about 10Å, in teal is the reconstruction which has clear density for the Cdc37 M/C, and in blue and maroon are the reconstructions with alternative conformations of Cdk4 N-lobe.

**Movie S4 Movie depicting profound conformational changes of Cdk4 and detailing the overall arrangement of the complex.**

